# Attenuated processing of vowels in the left hemisphere predicts speech-in-noise perception deficit in children with autism

**DOI:** 10.1101/2024.06.24.600191

**Authors:** Kirill A. Fadeev, Ilacai V. Romero Reyes, Dzerassa E. Goiaeva, Tatiana S. Obukhova, Tatiana M. Ovsiannikova, Andrey O. Prokofyev, Anna M. Rytikova, Artem Y. Novikov, Vladimir V. Kozunov, Tatiana A. Stroganova, Elena V. Orekhova

**Affiliations:** Center for Neurocognitive Research (MEG Center), Moscow State University of Psychology and Education, Moscow, Russian Federation; RSNO “Center for Curative Pedagogics”, Moscow, Russian Federation

**Author notes:** Corresponding author - Elena Orekhova. The first two authors contributed equally to the work.

**Keywords:** autism spectrum disorder (ASD), magnetoencephalography (MEG), speech-in-noise perception, vowels, formant structure, periodicity pitch, sustained processing negativity (SPN), children, auditory processing disorder

## Abstract

**Background:** Difficulties with speech-in-noise perception in autism spectrum disorders (ASD) may be associated with impaired analysis of speech sounds, such as vowels, which represent the fundamental phoneme constituents of human speech. Vowels elicit early (< 100 ms) sustained processing negativity (SPN) in the auditory cortex that reflects the detection of an acoustic pattern based on the presence of formant structure and/or periodic envelope information (*f0*) and its transformation into an auditory “object”.

**Methods:** We used magnetoencephalography (MEG) and individual brain models to investigate whether SPN is altered in children with ASD and whether this deficit is associated with impairment in their ability to perceive speech in the background of noise. MEG was recorded while boys with ASD and typically developing boys passively listened to sounds that differed in the presence/absence of *f0* periodicity and formant structure. Word-in-noise perception was assessed in the separate psychoacoustic experiment using stationary and amplitude modulated noise with varying signal-to-noise ratio.

**Results:** SPN was present in both groups with similarly early onset. In children with ASD, SPN associated with processing formant structure was reduced predominantly in the cortical areas lateral to and medial to the primary auditory cortex, starting at ∼ 150 - 200 ms after the stimulus onset. In the left hemisphere, this deficit correlated with impaired ability of children with ASD to recognize words in amplitude-modulated noise, but not in stationary noise

**Conclusions:** These results suggest that perceptual grouping of vowel formants into phonemes is impaired in children with ASD and that, in the left hemisphere, this deficit contributes to their difficulties with speech perception in fluctuating background noise.

## BACKGROUND

Autism spectrum disorder (ASD) is a group of neurodevelopmental conditions characterized by social and communication impairments and repetitive and restricted behaviors and interests [1]. Up to 70% of children with ASD have language delay, although the exact figures vary depending on the age group investigated and diagnostic criteria used [2–6]. Deficits in receptive and expressive language are associated with early ASD diagnosis [7] and worse outcome [4] and frequently observed even in verbal children with ASD [2,4]. While in many cases language deficits in ASD can be attributed to general level of cognitive development [8] and/or social motivation [4,9], there is a strong reason to believe that atypical processing of auditory information may contribute to the observed deficits [10–12].

A common difficulty faced by people with ASD, even those with normal or above normal IQ, is a poor listening ability under suboptimal acoustic conditions, such as background noise, both in experimental settings (for a review see [13]) and in real-life [14–17]. Several studies have linked the ability to perceive speech in noise to the fidelity of temporal processing in the brainstem [18–21]. While impaired temporal processing at the subcortical level in individuals with ASD [22–25] may affect their ability to perceive speech in noise, it is less clear whether processing of the basic phonetic properties of speech sounds in the auditory cortex contributes to this deficit. In this study, we address this question by examining in children with ASD the relationship between speech perception in noise and vowel processing in the auditory cortex.

Focusing on vowels may be interesting in two respects. First, vowels represent the simplest and ontogenetically and phylogenetically oldest phoneme constituents of human speech. They are the first speech sounds to be produced by human infants [26]. Vowel-like sounds are present even in the vocal repertoire of non-human primates [27]. Thus, atypical vowel processing may have serious effects on speech perception in noise and on language skills in general.

The second reason relates to the acoustic properties of vowel sounds. Vowels are *acoustic patterns* characterized by formant structure and common periodicity. Detection of acoustic patterns is a rapid and automatic process subserved by the auditory cortex [28–31]. The combination of formants, i.e., peaks in the frequency spectrum, determines the identity of the vowel, and the periodicity of the amplitude envelope defines its pitch (i.e., fundamental frequency, *f0*). Extraction of these complex features is followed by processing of the vowel as an auditory ’‘object” [32–34]. In the absence of linguistic context (i.e., when the vowel is not represented as part of a word), the spectral-temporal structure of the vowel remains the only auditory cue for the bottom-up grouping that governs its neural representation as a perceptually meaningful auditory object. There is some evidence that the ability to automatically group sound features is reduced in people with ASD [35–37], and it has been suggested that this deficit may contribute to their impaired speech perception in noisy environments, as the auditory system must rely on automatic grouping to effectively process speech [36]. However, whether automatic vowel processing in the auditory cortex in ASD is altered in a way that affects speech perception remains an open question.

The processing of auditory patterns and formation of auditory objects (figures) is associated with a sustained negative shift in neural current recorded with MEG/EEG [30,31,38–41], which we will refer to here as sustained processing negativity (SPN). It has been suggested that SPN reflects the persistent activity of “non-synchronized” neural populations [41,42]. These neurons are most abundant in non-primary auditory areas, are highly sensitive to complex sound features such as combinations of certain spectral and temporal parameters, and are thought to support representation of meaningful auditory patterns, including species-specific vocalizations or other ecologically relevant sounds [28,29,34,43–45]. The term “non-synchronized” refers to the property of these neurons to sustained firing for hundreds of milliseconds, especially when driven by their preferred continuous stimuli. These neurons are thought to encode temporally integrated spectral information [28,29,46] and transform it from an acoustic to a perceptual dimension [29]. In the case of vowels, this transformation seems to be necessary to form an integrated phonetic representation.

The presence of bottom-up grouping cues characterizing vowel sounds, such as periodicity of the amplitude envelope, formant structure, and especially the combination of these features, leads to a rapid (within the first 80 ms) and long-lasting (∼400 ms or longer) increase of the SPN in the auditory cortex [39,41,47]. The sensitivity of the SPN to bottom-up grouping signals and its ubiquitous presence in the neural response of children and adults suggests that it may serve as a candidate for detecting putative vowel processing dysfunction at the level of the auditory cortex in children with ASD.

Although previous EEG/MEG studies in ASD individuals have not examined vowel-induced SPN, many studies have investigated mismatch negativity/field (MMN/MMF) response to changes in vowels or syllables in the oddball paradigm. Despite considerable variation in results, their meta-analysis showed that individuals with ASD had increased MMN/MMF latencies and decreased MMN/MMF amplitudes in response to “different phoneme” deviations, but not to phoneme-duration or phoneme-pitch deviations [48], suggesting specific disturbance of phonetic processing. The MMN/MMF studies, however, had certain limitations. *First*, they have not identified auditory cortical areas involved in abnormal vowel processing in ASD, either due to limitations of sensor-level analysis or coarse localization of brain activity based on a template brain. *Second*, MMN/MMF studies typically focused on the peaks of event-related responses, perhaps missing differences in activation beyond these peaks. *Third*, for reasons still under debate, transient ERP/ERF peaks in children and adults differ in timing, polarity, and underlying neural processes [49–51], making it difficult to compare data across age groups. *Fourth*, since the auditory cortex is sensitive to any regular acoustic patterns [30,31,35], atypical MMN to vowels may reflect a deficit that is not specific to the processing of speech sounds.

In the present study, we aimed to clarify some of the previously unresolved questions regarding putative vowel processing deficit in ASD. *First*, we applied MEG imaging combined with individual brain models to localize cortical sources of atypical activity associated with analysis of vowels in children with ASD. *Second*, we tracked the entire timecourse of the vowel-related response in the auditory cortex rather than focusing on response maxima. *Third*, we focused on the SPN, which is present in both children and adults, overlapping with age-specific transient components of the auditory response, and may prove informative for testing putative vowel processing deficits in the developing brain. *Fourth*, to disentangle putative deficits related to the processing of formant structure (’‘speechness”) and periodicity of the vowel sound, we used synthetic vowels in which the formant structure and *f0* were either preserved or modified so that they could be analyzed separately.

We next asked whether atypical vowel processing is associated with impairment in the ability of children with ASD to distinguish speech from background noise. Such a possibility is worth investigating given the psychoacoustic evidence for the importance of vowels for speech intelligibility [52] and the evidence for impaired speech-in-noise recognition in ASD individuals, including those with normal-range IQ and audiometrically normal hearing (for a review see [13]).

Although both words/syllables and sentences have previously been used to test speech-in-noise perception deficit in children with ASD, in the present study we decided to apply the “words-in-noise” (WiN) paradigm. Words are less taxing on working memory than sentences, and their repetition is less dependent on cognitive ability, allowing this test to be used for verbal children with below-average intelligence. Our recent study showed that the ability to recognize two-syllable high-frequency words in background noise in verbal children with ASD does not correlate with their IQ [53]. Unlike the recognition of sentences, which carry higher-order linguistic information (overall tonal contour, stress pattern, syntactic structure, etc.), the recognition of isolated words relies more on bottom-up grouping cues. Thus, the words-in-noise paradigm is better suited than the sentences-in-noise paradigm to investigate the perceptual effects of a possible dysfunction in the relatively low-level cortical grouping processes underlying vowel identification.

To compare WiN perception in children with ASD and TD children, we used two types of background noises: stationary (ST) and amplitude-modulated (AM). Dips (i.e., short periods with more favorable SNR) in the AM noise allow listeners to improve speech recognition. This improvement, referred to as masking release, is attributed to the listener’s ability to detect the target speech signal during the dips [54]. It has been suggested that atypically low masking release in individuals with ASD is due to a reduced ability to integrate fragments of auditory information into meaningful words or sentences [55,56]. However, deficit in phonetic processing, especially of vowels, may also play a role, as the preserved spectral-temporal structure of vowels is particularly important for speech perception when speech is immersed in AM noise [52,54].

To summarize, we hypothesized that processing of natural auditory objects - vowels - is atypical in children with ASD and that this deficit may contribute to their degraded perception of words in noise. To test this hypothesis, we recruited verbal children with ASD and TD children and examined group differences in SPN using magnetoencephalography (MEG). To find out which vowel features processing is affected in ASD, we used synthetic periodic and aperiodic vowels, as well as complex periodic sounds that lacked vowel formant structure. We aimed to investigate when (in terms of processing timing) and where (in terms of auditory cortical regions) the timecourse of SPN in children with ASD diverges from the typical activation pattern. We then tested the relationship between atypical SPN activation patterns in children with ASD and their word recognition abilities in the ST and AM noise. Given the importance of vowels for ’‘dip listening”, we hypothesized that vowel processing deficit in children with ASD, if present, should predominantly affect their performance in AM noise.

## MATERIALS AND METHODS

### Participants

The study included 39 boys with ASD aged 6.9 – 13.0 years and 35 typically developing (TD) boys of the same age range. The TD children from the control group were recruited through advertisements in the media. None of the TD participants had known neurological or psychiatric disorders. Twenty two of 39 TD participants were the same as in our recent developmental study devoted to comparison of sustained negative responses to periodicity and formant composition of vowels in children and adults [41]. The participants with ASD were recruited through several sources (advertisements in the media, consulting centers, an educational center affiliated with the Moscow State University of Psychology and Education).

The ASD diagnosis was confirmed by an experienced psychiatrist who was the author of the current study (N.A.Y.) and was based on the Diagnostic and Statistical Manual of Mental Disorder (5th ed.) criteria as well as an interview with the child and parents/caregivers. In addition, parents of all children were asked to complete the Russian translation of parental questionnaires: Social Responsiveness Scale for children [57] and Social Communication Questionnaire (SCQ-Lifetime) [58] and most of them completed these questionnaires (Table 1). Intellectual abilities were assessed using the KABC-II test, and the Mental Processing Index (MPI) was used as an IQ equivalent [59]. This index excludes tasks that require verbal concepts, verbal reasoning, and cultural knowledge and is recommended for evaluation of non-verbal abilities in both TD children and those with ASD [60].

**Table 1.** Characteristics of the samples.

| | Mean (range) ± SD or Median (range) | | Student T-test /<br>M-W U test*, Z | p |
| --- | --- | --- | --- | --- |
|  | *TD* | *ASD* |  |  |
| **Age** | Mean = $9.8 \pm 1.8$<br>(7.3 - 13.0)<br>N = 39 | Mean = $10.2 \pm 1.7$<br>(6.9 - 12.8)<br>N = 35 | T = 0.87 | 0.39 |
| **MPI IQ**<br>**(KABC-II)** | Mean = $116.2 \pm 13.3$<br>(85 - 138)<br>N = 39 | Mean = $83.1 \pm 15.8$<br>(54 - 128)<br>N = 34 | T = 9.7 | < 0.0001 |
| **SRS** | Mean = $37.6 \pm 20.0$<br>(3 - 84)<br>N = 37 | Mean = $99.5 \pm 27.2$<br>(33 - 157)<br>N = 34 | T = 11.0 | < 0.0001 |
| **SCQ** | Mean = $6.3 \pm 3.2$<br>(2 - 16)<br>N = 38 | Mean = $24.8 \pm 6.4$<br>(12 - 35)<br>N = 32 | T = 15.6 | < 0.0001 |
| **Language**<br>**total**<br>**score\*\*** | Median = 94.9<br>(86.3 - 98.6)<br>N = 29 | Median = 79.7<br>(35.4 - 94.1)<br>N = 32 | Z = 6.2 | < 0.0001 |

N - number of participants for whom information is available.
\* Mann-Whitney U Test was applied for the group comparison in Language total scores, because their distribution in the ASD group was significantly different from normal.
\*\* language proficiency was estimated using the RuCLAB test.

Hearing status of all participants was determined using pure tone air conduction audiometry with a AA-02 clinical audiometer (“Biomedilen”). Auditory sensitivity was tested at 500, 1000, 2000, 4000 Hz, and the threshold averaged over the four frequencies was calculated separately for each ear. All participants had normal hearing (threshold< 20 dB HL) in either ear [61].

The Ethical Committee of the Moscow State University of Psychology and Education approved this investigation. All children gave verbal consent to participate in the study and their caregivers gave written consent to participate.

### Assessment of the general level of language development

The language test incorporated 12 subtests from the Russian Child Language Assessment Battery, RuCLAB [62], which evaluates expressive and receptive skills in vocabulary (production and comprehension of word), morphosyntax (sentence production and comprehension), and discourse (text production and comprehension). The audio material was provided by a professional female native Russian speaker and recorded in studio conditions. All test stimuli were delivered through the AutoRAT application [63]. The tests took place in a quiet and child-friendly environment. Before each subtest, participants received instructions and performed 2-3 practice trials, which were not included in the final analysis. The sequence of the test items was the same for all participants. During testing, the examiner completed a paper protocol. Additionally, the sessions were videotaped, adhering to ethical standards. The description of the test components and details on the scoring are provided in the Supplementary Methods and Table S1. To characterize the overall level of language proficiency, the arithmetic mean of scores on all subtests was calculated. This mean score is hereinafter referred to as the “total language score”.

### Words-in-Noise (WiN) test

#### Stimuli

The Words-in-Noise (WiN) test includes 160 two-syllable lemmatized Russian nouns with high imagery (ability to evoke mental images). These words were frequently occurring words according to the List of Frequent Nouns [64] and/or the list of 300 words most frequently used in daily life [65], and corresponded to the age norm for children 6 years and older. The words were spoken by a 35-year-old woman with neutral, unemotional intonation and were recorded with studio recording equipment. The mean duration of the words was 694 ms (SD = 77 ms).

Mean loudness of each word was adjusted to correspond to that of 45 dB SPL pink noise using the “spl’’ function of a third-party software package for MATLAB R2020a [66]. The masking noise was of two types: stationary pink noise (ST) and amplitude modulated pink noise (AM) and was synthesized using ’‘pinknoise” and ’‘ammod” functions of MATLAB R2020a (MathWorks, Inc.). AM noise was obtained by modulating ST noise with a 10 Hz sinusoidal function (i.e., pink noise was interrupted 10 times per second); modulation started at the 0° phase. Power spectral density of the masking noise decreases proportional to its frequency (1/f), 3 dB per octave. Noise lasted for 1 second and started 75 ms before word onset. The rise/fall of each 1-second noise signal was smoothed at 1 ms intervals. Words were presented at a fixed sound pressure level. The sound pressure level of the masking noise was varied at four signal-to-noise ratios (SNR): -0, -3, -6, and -9 dB.

#### Testing procedure

The stimuli were presented through Sony WH-XB900N headphones with the noise-canceling function off. The headphones were calibrated using a CEM DT-815 sound level meter. A stationary pink noise of 45 dB was taken as the reference. The experiment began with a training session during which 10 words were presented randomly against a background of ST or AM noise (3 to -3 SNR). The participant was asked to repeat the word after each presentation. Only the exact repetition was considered a correct answer. The training lasted until the child internalized the instruction, but no more than 10 min. If the instruction was successfully learned, the experimenter proceeded to the main part of the test, which included four blocks in the sequence of 0, -3, -6, and -9 dB SNR, where the 0 SNR condition was the easiest and the -9 SNR condition was the most difficult. Each block contained 40 words - 20 words for each noise type (ST and AM). The type of the masking noise varied within a block in a pseudorandomized order (no more than three consecutive presentations of the same noise type). Words were selected randomly from the list and presented in one of the conditions. Each word was presented only once. For each condition (4 SNR levels, 2 types of masking noise), the number of correctly recognized words was counted. At the end of the main part of the experiment, words that the participant did not repeat correctly were presented without noise. All of the children in this study were able to repeat the words presented to them without noise. The WiN test was presented to 30 (of 39) TD children and to 29 (of 35) children with ASD. All TD and 27 of 29 ASD participants were able to complete the test. The full data (MEG, results of WiN test, MPI IQ scores) were available for 26 participants. Therefore, partial correlation analyses were conducted on this smaller sample of participants with ASD.

#### Scoring

Although the words corresponded to the developmental level of the participants, some of them were particularly easy or difficult to recognize. We excluded from the analysis 18 words that were ’‘easiest” or “most difficult” according to the accepted criteria (see Supplementary Methods). For each child, we then calculated the average percent of correct responses in each of the SNRs (0dB, -3dB, -6dB, -9dB) in the ST and AM noise conditions. Many participants of both groups (14 of 30 TD and 14 of 27 ASD) gave 0% correct responses in the most difficult -9 dB ST noise condition. Therefore, we excluded the -9 dB SNR condition from further analysis. For each child, we then calculated the average percent of correct responses for each SNR level (0dB, -3dB, -6dB) in the ST and AM noise conditions (WiNst and WiNam scores, respectively).

### MEG experiment

#### Stimuli

The experimental paradigm used in the present study is identical to that described in Orekhova et al. [41]. We used the four types of synthetic vowel-like stimuli previously used by Uppenkamp et al. [67] and downloaded from ’‘http:/ medi.uni-oldenburg.de/members/stefan/phonology_1/”. Five strong vowels were used: /a/ (caw, dawn), /e/ (ate, bait), /i/ (beat, peel), /o/ (coat, wrote) and /u/ (boot, pool).

The synthetic periodic vowels consisted of damped sinusoids, which were repeated with a period of 12 ms, so that the fundamental frequency of each vowel was 83.3 Hz. The carrier frequencies of each vowel were kept fixed at the four lower formant frequencies, which were chosen in a typical range of an adult male speaker. These are further referred to as periodic vowels. These regular vowel stimuli have been modified, as described below, to generate three other classes of stimuli: non-periodic vowels, periodic non-vowels, and non-periodic non-vowels. To violate periodicity, the start time of each damped sinusoid was jittered within ±6 ms relative to its start time in the original vowel, separately for each formant. Despite the degraded voice quality (hoarse voice), these non-periodic sounds were perceived as vowels. To violate formant constancy, the carrier frequency of each subsequent damped sinusoid was randomly chosen from a set of eight different formant frequencies used to produce regular vowels and randomized separately for each formant (Frequency range: formant 1 = 270 - 1300 Hz; formant 2 = 850 - 2260 Hz; formant 3 = 1750 - 3000 Hz; formant 4 = 3300 - 5500 Hz). Both periodic and non-periodic sounds with disrupted formant structure were perceived as noises rather than vowels (i.e. non-vowels). The following four stimulus types were presented during the experiment: (1) periodic vowels (/a/, /i/, /o/); (2) nonperiodic vowels (/a/, /u/, /e); (3) three variants of periodic non-vowels and (4) three variants of non-periodic non-vowels. The spectral composition of these stimuli is given in Supplementary Figure S1 (see also [41]).

Two hundred seventy stimuli of each of the four classes were presented, with three stimulus variants equally represented within each class (N = 90). All stimuli were presented in random order. Each stimulus lasted 812 ms, including rise/fall times of 10 ms each. The interstimulus intervals (ISIs) were randomly chosen from a range of 500 to 800 ms.

The non-periodic non-vowels were used as control stimuli. The contrasts of interest were (1) “non-periodic vowels versus non-periodic non-vowels”, (2) “periodic non-vowels versus non-periodic non-vowels” and (3) “periodic vowels versus non-periodic non-vowels”. By comparing these contrasts in the ASD and TD groups we investigated group differences in the processing of formant structure, periodicity/pitch, or a combination of these features in a natural vowel.

#### Procedure

Participants were instructed to watch a silent video (movie/cartoon) of their choice and ignore the auditory stimuli. Stimuli were delivered binaurally via plastic ear tubes inserted into the ear channels. The tubes were attached to the MEG helmet to prevent possible noise from their contact with the subject’s clothing. The intensity was set at 90 dB SPL. The experiment included three blocks of 360 trials, each block lasting around 9 minutes with short breaks between blocks. If necessary, the parent/caregiver remained with the child in the MEG shielded room during the recording session.

### MRI data acquisition and processing

In all participants with ASD and in 28 TD participants T1-weighted 3D-MPRAGE structural image was acquired on a Siemens Magnetom Verio 3T scanner (Siemens Medical Systems, Erlangen, Germany) using the following parameters: [TR 1780 ms, TE 2.78 ms, TI 900 ms, FA 9°, FOV 256 × 256 mm, matrix 320 × 320, 0.8 mm isotropic voxels, 224 sagittal slices]. In 9 TD subjects MRIs were acquired at a 1.5T Philips Intera. In 2 TD subjects 1.5T GE Brivo MR355/MR360 was used. Cortical reconstructions and parcellations were generated using FreeSurfer v.7.4.1 [68,69].

### MEG data acquisition, preprocessing and source localization

MEG data were recorded at the Moscow Center for Neuro-cognitive Research (MEG-Center) using Elekta VectorView Neuromag 306-channel MEG detector array (Helsinki, Finland) with 0.1 - 330 Hz filters and 1000 Hz sampling frequency. Bad channels were visually detected and labeled, after which the signal was preprocessed with MaxFilter software (v.2.2) in order to reduce external noise using the temporal signal-space separation method (tSSS) and to compensate for head movements by repositioning the head in each time point to an “optimal” common position (head origin). This position was chosen individually for each participant as the one that yielded the smallest average shift across all data epochs after motion correction.

Further preprocessing steps were performed using MNE-Python software (v.1.4.1) [70]. The data were filtered using notch-filter at 50 and 100 Hz and a 110 Hz low-pass filter. Periods in which peak-to-peak signal amplitude exceeded the thresholds of 7e-12 T for magnetometers or 7e-10 T/m for gradiometers, and “flat” segments in which signal amplitude was below 1e-15 T for magnetometers or 1e-13 T/m for gradiometers were automatically excluded from further processing. To correct cardiac and eye movement artifacts we recorded ECG, vEOG, and hEOG and used a signal-space projection (SSP) method. Next, we excluded from analysis data segments in which head rotation exceeded a threshold of 10 degrees/s along one of the three space axes, head velocity exceeded a threshold value of 4 cm/s in 3D space, or head position deviated from the origin position by more than 10 mm in 3D space. The data were then epoched from -0.2 s to 1 s relative to stimulus onset. The mean number of artifact-free data epochs was initially higher in the TD participants (TD: 997 vs ASD: 927, p < 0.05). To equalize this number, we randomly removed 70 epochs for each TD participant. The resulting mean number of clean epochs for each subject and stimulus type was 231 (range 141 - 340) and 231 (range 131 - 347) for TD and ASD children, respectively. The epoched data were averaged separately for the four experimental conditions and then baseline corrected by subtracting the mean amplitude in -200 - 0 ms prestimulus interval.

To obtain the source model, the cortical surfaces reconstructed with the Freesurfer were triangulated using dense meshes with about 130,000 vertices in each hemisphere. The cortical mesh was then resampled to a grid of 4098 vertices per hemisphere, corresponding to a distance of about 4.9 mm between adjacent source points on the cortical surface.

To compute the forward solution, we used a single layer boundary element model (inner skull). Source reconstruction of the event-related fields was performed using the standardized low-resolution brain electromagnetic tomography (sLORETA) [71]. Noise covariance was estimated in the time interval from -200 to 0 ms relative to stimulus onset. To facilitate comparison between subjects, the individual sLORETA results were morphed to the fsaverage template brain provided by FreeSurfer.

### MEG data analysis

#### Data analytic plan

*First*, to find out whether SPN associated with processing vowel periodicity and formant structure presents in both TD children and children with ASD, we compared responses to test stimuli (periodic vowels, non-periodic vowels, periodic non-vowels) and control stimuli (non-periodic non-vowels) separately in TD and ASD groups. This was done in the sensor space using a global root mean square RMS signal and then in the source space using the sLORETA values in combination with the nonparametric permutation test with spatiotemporal threshold-free cluster enhancement (TFCE) [72]. The TFCE analysis was performed in the left and right regions of interest (ROIs) broadly overlapping the auditory cortex. To analyze temporal characteristics of the responses associated with processing of periodicity or formant structure, we compared timecourses of neural currents evoked by test and control stimuli in those sources where the effects identified by TFCE analysis were most significant.

*Second*, we used TFCE cluster analysis in the ROIs to test for the group differences in differential responses (e.g. response to periodic vowel minus response to control stimulus). Then, we analyzed the timecourses of the group differences in the “most significant” sources identified by TFCE analysis.

*Third*, when group differences in differential responses were found, we tested for their correlation with WiN scores in children with ASD using partial correlation analysis to account for the effects of age and IQ.

#### Root mean square (RMS) analysis

For the sensor level analysis, all subjects’ data were transformed to a common standard head position (to [0, 0, 45] mm, default MaxFilter parameter). RMS metric was computed based on the signal from all gradiometer sensors and compared between test (periodic vowels, periodic non-vowels, non-periodic vowels) and control (non-periodic non-vowel) conditions point-by-point in the 0 - 800 ms stimulation interval using paired t-tests.

#### Spatiotemporal cluster analysis

The spatiotemporal clustering analysis was performed in the ROIs that were selected in the fsaverage template brain so as to broadly overlap the left and right auditory cortex and nearby areas (Figure 3). These ROIs were identical to those used in our previous study [41]. The direction of the source current in the ROIs was aligned using the MNE-python function ’‘label_sign_flip”. We then verified that in each participant and hemisphere, the timecourse averaged over all the point sources and across conditions had a negative sign between 300-800 ms, consistent with the sustained negativity observed in response to these stimuli in the auditory cortex [39,41] and a positive sign of the P100m component between 50-150 ms. The source current data were cropped to 0-800 ms timeregion and downsampled to 500 Hz.

TFCE spatiotemporal cluster analysis was used to compare evoked source currents between groups. The TFCE procedure derives the spatiotemporal cluster-level statistics for each data point *p* by using a weighted average between the cluster extent *e* (i.e., number of connected above-threshold data points) and the cluster height ℎ (i.e., the statistical value of the data point), calculated according to the formula:

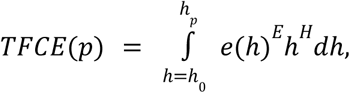

where default values of *E* = 0. 5 and *H* = 2 were applied [72], and starting threshold ℎ_0_ = 0, step size *dh* = 0. 4 were set up for the better approximation. Permutation tests with TFCE effectively address the issue of multiple comparisons, and in this study, we computed 5000 TFCE permutations. For each data point, a corrected p-value was calculated by determining the proportion of permutations in which the TFCE output was greater than or equal to the original TFCE output. The null hypothesis of no cluster of the difference between the groups or conditions (within each group) was rejected at p < 0.05. As no interhemispheric spatial-adjacency-based clusters were expected, the ’‘check_disjoint=True” option was applied, which is equivalent to running the test separately for each hemisphere.

To identify clusters showing significant differences between test and control stimuli we used one-sample permutation test with spatiotemporal TFCE (’‘stats.permutation_cluster_1samp_test” MNE Python function) separately in each group. In this test, the inputs were the differences between responses to test and control stimuli that were permuted within a single subject.

To identify clusters of significant Group × Condition interactions, we employed a two-sample permutation test with spatiotemporal TFCE (’‘stats.spatio_temporal_cluster_test” MNE Python function), where the inputs were the differences between responses to the test and control stimuli for each subject from the ASD and TD groups.

When clusters of significant differences were found, we examined timecourses of activity in the “most significant” sources within these clusters. In this case, we did not consider sources beyond the “STG+” region (areas A1, 52, LBelt, PBelt, MBelt, RI, A4, TA2, PI and PoI1, according to the HCPMMP1 atlas [73]). All the STG+ areas are structurally connected [76,77] and participate in processing of auditory information [78–80]. On the other hand, activity observed in response to auditory stimuli in superior insula and superior temporal sulcus may reflect point spread from auditory cortical regions [74,75] and inspection of the polarity of the current induced by auditory stimuli in our study confirms this (Figure S3).

To select the “most significant” sources for timecourse analysis, we applied an approach similar to that used in [81]. Specifically, we selected sources within the cluster with the average p-values that were below the global average within this cluster. The sources were defined separately for the left and right hemispheres and for each comparison performed (i.e., test-vs-control and ASD-vs-TD contrasts).

To analyze correlations of the SPN with WiN scores in children with ASD, the activity of the “most significant” sources was averaged in the time interval that displayed strongest differences between TD and ASD groups. The temporal boundaries of this interval were restricted to the mean onset and end times of significant group differences in the “most significant” sources (see Supplementary Figure 2). Using linear regression analysis, we then calculated the adjusted SPN (SPNadj) by regressing from the response to the test stimulus the magnitude of the response to the control (non-periodic non-vowels) stimulus and the square root of the mean number of averaged epochs per condition. Thus, by computing SPNadj, we accounted for large interindividual variability in the magnitude of auditory responses and for differences in the number of averaged epochs.

### Statistical analysis

Nonparametric tests (Mann-Whitney U Test, Spearman correlations, or partial Spearman correlations) were used when the distribution differed significantly from normal according to Shapiro-Wilk’s W test (p < 0.05). Otherwise parametric tests (T-test, Pearson correlations, Pearson partial correlations) were used. To calculate partial correlations we used the ’‘pcor” function in R. Group or condition-related differences in neural responses were assessed using TFCE cluster analysis (see above). In case of multiple comparisons, the False discovery rate (FDR) method of Benjamini and Hochberg [82] for correcting p-values was applied. The accepted significance level was p < 0.05. To analyze the effect of SNR (0 dB, -3 dB and -6 dB) and type of masking noise (ST, AM) on the difference in WiN scores between the TD and ASD participants, we employed Mann-Whitney U Test and/or linear mixed model analysis using the ’‘lmer” function in R.

## RESULTS

### Characteristics of the samples

The characteristics of the samples are summarized in Table 1. Participants with ASD compared to TD participants had significantly higher SRS and SCQ scores. In the case of the SRS questionnaire, all but 3 of 34 tested subjects with ASD diagnosis had total scores above 60 points cut-off for ASD. For the SCQ questionnaire, all but 3 of 32 tested ASD subjects scored above the 15 points cut-off. The three ASD subjects who scored below the cut-off on the SRS questionnaire scored above the cut-off on the SCQ questionnaire, and vice versa. In the TD sample all but 5 children scored below ASD threshold on the SRS questionnaire, and all but one scored below ASD cut-off on the SCQ questionnaire. Given the sensitivity and specificity of the SRS and SCQ questionnaires [83,84], these results suggest that our samples of ASD and TD children are very well separated on autistic traits.

The participants with ASD as compared with TD participants had significantly lower MPI IQ scores. 18 ASD subjects (53%) had the MPI IQ scores below 85, which corresponds to “below-average” intelligence [59]. This percentage of ASD children with below-average intelligence roughly corresponds to that reported in 8-years-olds with ASD in USA in year 2020 (61% [85]) and is slightly lower than that reported in Chinese children born 2002-2008 and aged 6-12 years (∼64% [86]).

All, but two children with ASD displayed a history of language delay or impairment, defined by parents’ reports of lack of two-word combinations at age three or the presence of language problems at the time of the diagnostic assessment. Six children with ASD experienced language regression at some point in their development.

### Words in noise (WiN) test results in TD and ASD groups

Children with ASD recognized significantly fewer words than TD children in AM noise at all SNRs and in ST noise at -3 and -6 dB SNR (Figure 1A). Both TD and ASD children performed better in case of the AM than ST noise, with the exception of the 0 dB SNR condition in the ASD group, when no ’‘masking release” (i.e., improved performance in AM compared to ST-noise) was observed (Figure 1B).

**Figure 1.**
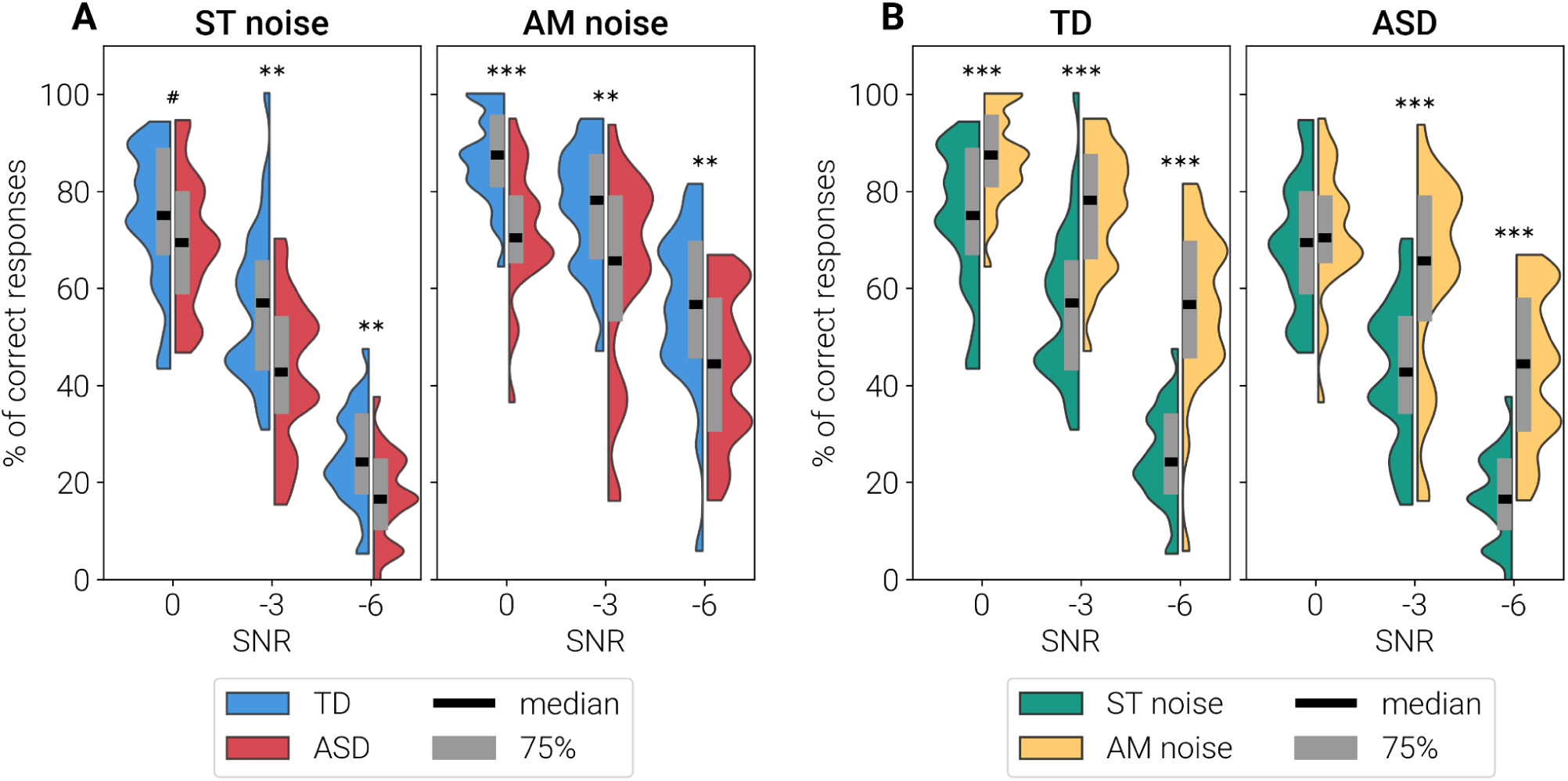
Mean percentage of correctly repeated words presented against a background of masking noise. **A:** Comparison of performance in TD and ASD groups, separately for the stationary and amplitude-modulated noise conditions. **B:** Comparison of performance in stationary and amplitude-modulated noise, separately for TD children and children with ASD. # p < 0.1, ** p < 0.01, *** p < 0.001 (Mann-Whitney U Test).

Linear mixed model with main effects of Group, noise Type (ST, AM) and noise Level (0, -3 and -6 dB SNR) and random intercept for subject have found significant interaction between Group and Type (estimate = -0.10, SE=0.046, t(270)=-2.21, p=0.028). Presence of gaps in noise resulted in lower masking release in the ASD group than in TD group. Given this effect, as well as the previous literature that show different masking properties of the ST and AM noise [55,56], for each child we calculated WiNst and WiNam scores for ST and AM noise separately, as the percentage of correct responses averaged over SNR levels.

In both groups, WiNst and WiNam scores improved with age (Spearman correlations; WiNst: TD, R(N=30) = 0.43, p = 0.02; ASD R(N=26) = 0.50, p = 0.009; WiNam: TD R(N=30) = 0.56, p = 0.001, ASD R(N=26) = 0.51, p = 0.008). WiN scores did not correlate with IQ in either group (Spearman correlations; WiNst: TD R(N=30) = 0.08, p = 0.65, ASD R(N=26) = -0.27, p = 0.18; WiNam: TD R(N=30) = 0.10, p = 0.58, ASD R(N=26) = 0.02, p = 0.93).

To sum up, the ability to recognize words in noise was impaired in children with ASD compared to their TD peers, was independent of IQ, and improved with age.

### WiN performance and general language abilities in TD and ASD groups

As expected, children with ASD had significantly lower total language scores than TD participants (Mann-Whitney U Test: Ntd = 29, Nasd = 32, Z = 6.2, p < 0.0001, η2 = 0.64). The scores improved with age in control participants (Spearman R (N=29) = 0.74, p < 0.0001), but not in children with ASD (Spearman R (N=32) = 0.21, p = 0.23). Unlike WiN scores, language total scores correlated with IQ (Spearman correlations, TD (N=29): R = 0.35, p = 0.06; ASD (N=31): R = 0.46, p = 0.008).

### MEG results: sensor-level analysis

Figure 2 shows the grand average auditory evoked field waveforms expressed as root mean square (RMS) signals calculated over all 204 gradiometer channels. Compared to control stimuli, stimuli characterized by temporal regularity (periodic non-vowels), formant structure (non-periodic vowels), or a combination of these features (periodic vowels) caused a transient decrease in RMS around 100 ms relative to post stimulus onset (time range of the child P100m component), followed by its prolonged increase after ∼150 ms that lasted up to 400 ms or longer. The RMS amplitude provides a measure of the magnetic field strength across the MEG sensors regardless of polarity, and is blind to the direction of the condition-related difference in the cortical currents [87]. In our previous study we have shown that both the decreases and increases in RMS in response to vowel-like versus control stimuli were explained by sustained negative shift of current in neural activity of the superior temporal cortex [41], i.e., SPN. To account for current polarity, we further analyzed the data at the cortical source level.

**Figure 2.**
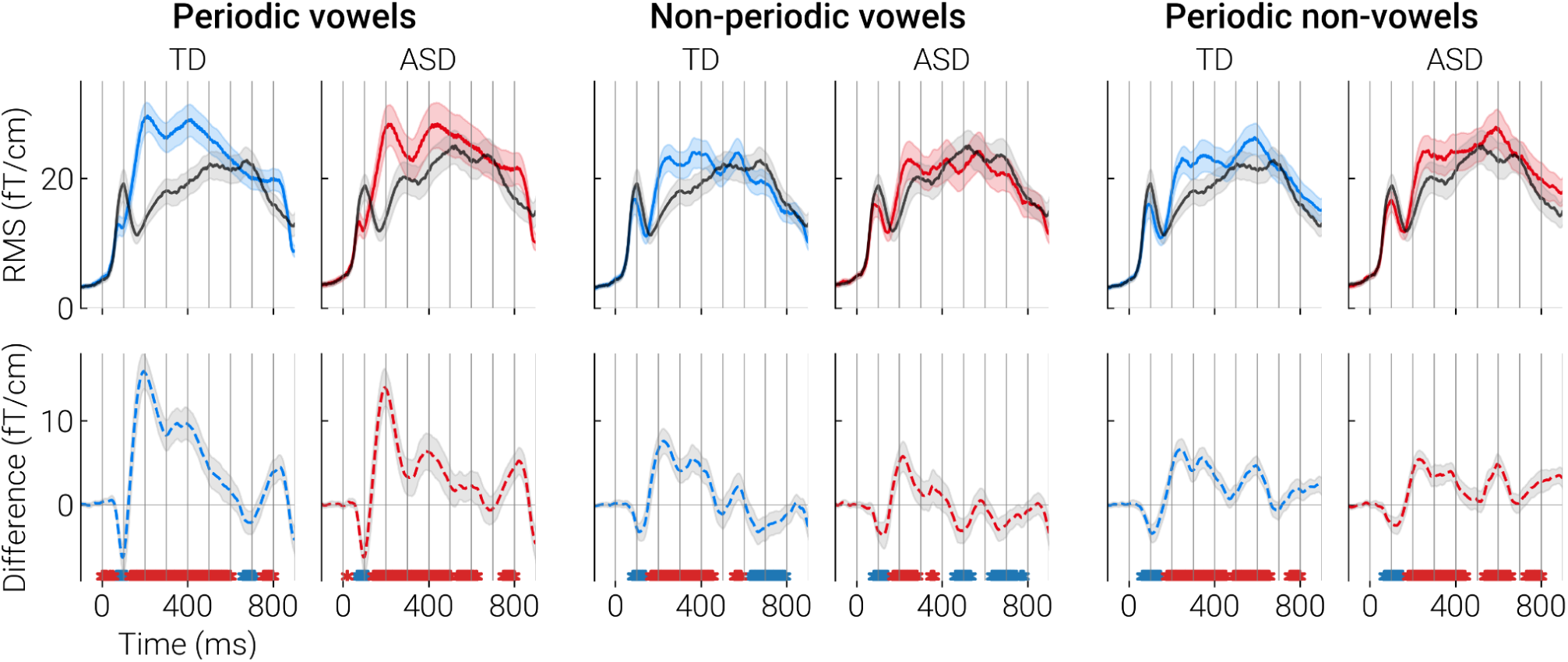
Grand average RMS response waveforms and the RMS difference waves in ASD and TD groups. Zero point at the horizontal axis corresponds to the onset of 800-ms stimulus. RMS was calculated over all gradiometer channels. Black line denotes control condition (non-periodic non-vowels) and colored lines denote test conditions (periodic vowels, non-periodic vowels, periodic non-vowels). Dashed lines indicate differential responses. The asterisks under the dashed lines correspond to significant point-by-point differences between the test and control conditions: red color - control < test; blue color - control > test (paired t-test, p < 0.05, FDR corrected). The shaded areas mark 95% confidence intervals.

### MEG results: analysis of the source current

#### Effects of periodicity and formant structure in TD and ASD participants

To test whether children with ASD, similarly to TD children [41], respond to patterned acoustic input with a SPN in the auditory cortical areas, we applied spatiotemporal cluster analysis separately in TD and ASD groups. SPN was defined as an increase in negativity in the evoked source current for the Test versus Control contrasts, i.e., a negative sign of the difference between vowel or complex periodic sounds (periodic non-vowels) and a control non-periodic non-vowel sound.

Cluster analysis was performed in the ROIs broadly overlapping the auditory cortex and nearby regions where auditory evoked activity was observed (Figure 3). Before cluster analysis, the direction of dipole sources within the ROIs was adjusted to correspond to the dominant direction of source current in the auditory cortex (see Methods for details). In both groups, periodicity, formant structure, and their combination were associated with bilateral clusters of negative differential responses, i.e., SPN that lasted several hundred milliseconds (Figure 3). SPN clusters encompassed primary and secondary auditory cortex and adjacent areas. In case of non-periodic vowels, the cluster of negative differences (test > control) was followed by (or co-existed with) a cluster of positive differences (test < control). In our previous study a similar pattern of differences between non-periodic vowel and control stimuli was observed in neurotypical individuals (see [41] for discussion). The temporal evolutions of the clusters of differences between test and control stimuli in the TD and ASD groups are shown in Supplementary videos S1-S6.

**Figure 3.**
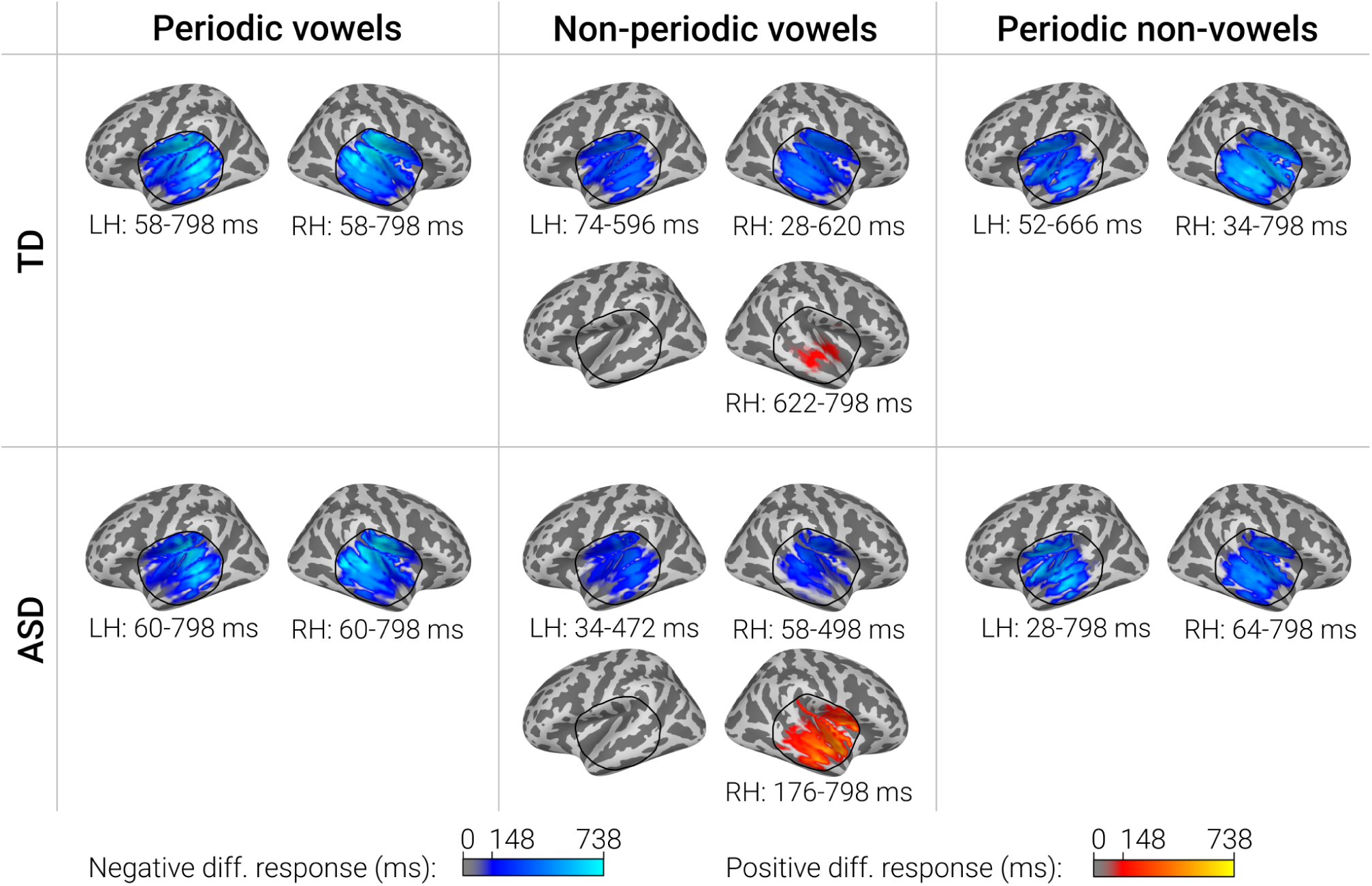
Significant clusters of the differences in the evoked source current between test (periodic vowel, non-periodic vowel, periodic non-vowel) and control (nonperiodic non-vowel) conditions in TD and ASD groups. Blue colors correspond to SPN, i.e. a more negative source current to the test compared to the control condition, red colors - to the opposite direction of the difference. Color intensity indicates the duration of the cluster. Black line indicates the border of the region used for cluster analysis. Note that the 798 ms point is the last time point analyzed.

Figure 4 shows the sources in which the differences between periodic vowels and control stimuli identified by cluster analysis were most significant (Figure 4 A, B), and the mean timecourses in these sources (Figure 4 C, D). The respective differential responses (periodic vowels minus control stimuli) and their significance (i.e. difference from zero) are shown in Figure 4 E, F and Figure 4 G, H. Similar illustrations for non-periodic vowels and periodic non-vowels are given in the Supplementary materials (Figures S4 and S5).

**Figure 4.**
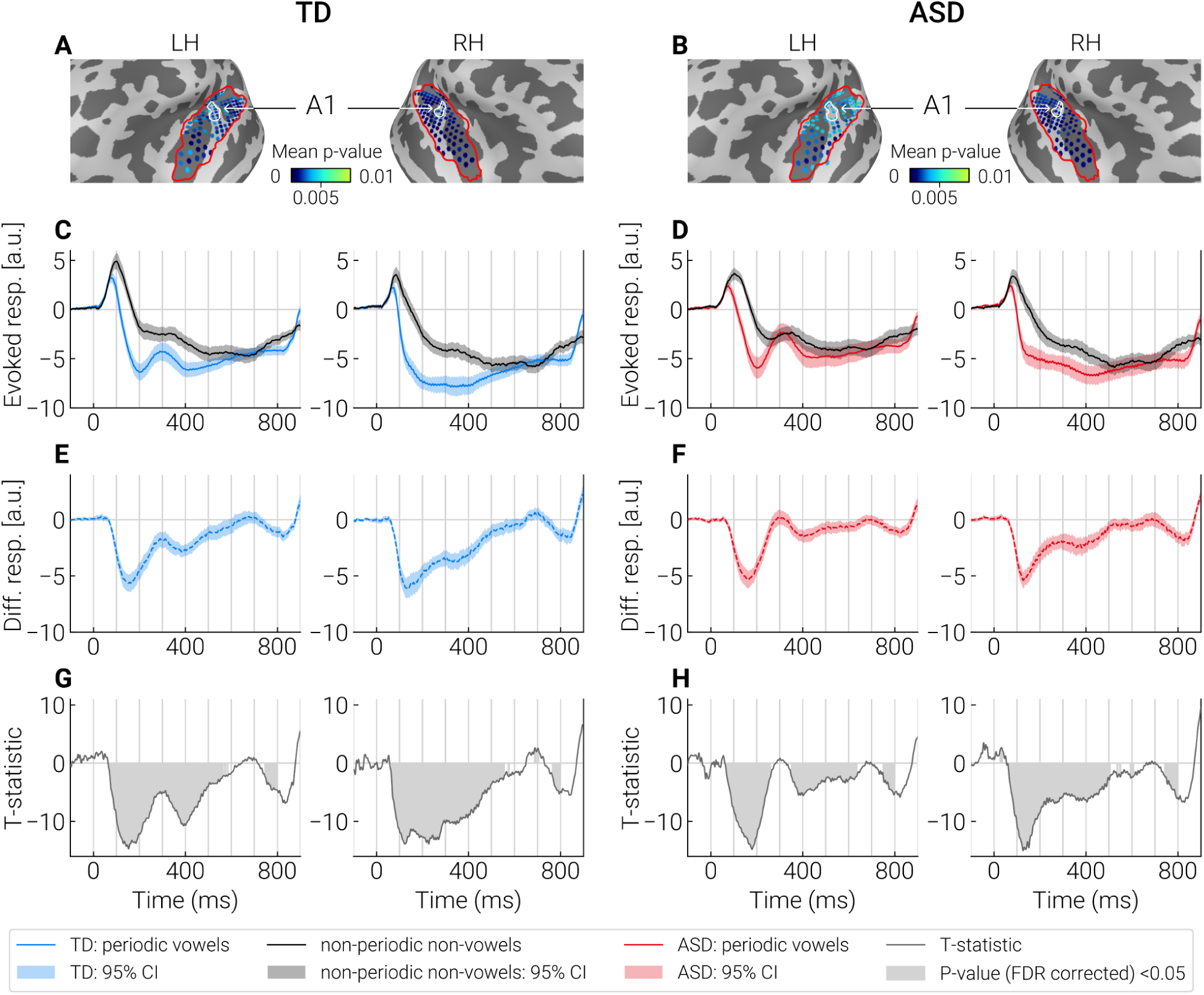
Comparison of evoked responses to the periodic vowels and control stimuli (non-periodic non-vowels) in the “most significant” dipole sources identified by cluster analysis. **A, B:** The “most significant” dipole sources within STG+ region (outlined with a red contour) are marked by blue dots. Color shade (light blue to dark blue) indicates significance of the differences between test and control conditions in the respective point sources. The primary auditory cortex (A1) is outlined with a white contour. **C, D:** Averaged neural current timecourses in the “most significant” sources. **E, F:** The difference between timecourses of current evoked by the test and control stimuli. **G, H:** T-statistics reflecting a pointwise comparison of the response timecourses to test and control stimuli. Significant differences (p < 0.05, FDR corrected) are marked in gray. Supplementary figures S4 and S5 show the similar results for non-periodic vowels and periodic non-vowels.

In both groups and in response to all test stimuli, significant SPN (i.e. negative sign of the difference in the source current between responses to test and control stimuli) began earlier than 100 ms after stimulus onset and then persisted for approximately 500 ms. This timing is consistent with previous results indicating early neural discrimination of auditory patterns characterized by grouping cues [30,88], which in our study were represented by frequency composition and/or periodicity. The group differences in the SPN are described in the next section.

#### Group differences in SPN related to the processing of periodicity and formant structure

TFCE cluster analysis yielded bilateral clusters of the group differences for differential responses (test – control condition) for non-periodic and periodic vowels (i.e., sounds characterized by the presence of formant structure), but not for periodic non-vowels. The spatial location of the clusters on the surface of the ’‘inflated” brain is shown in Figure 5. Supplementary Figure 6 shows the same clusters projected onto the surface of white matter for a three-dimensional representation.

**Figure 5.**
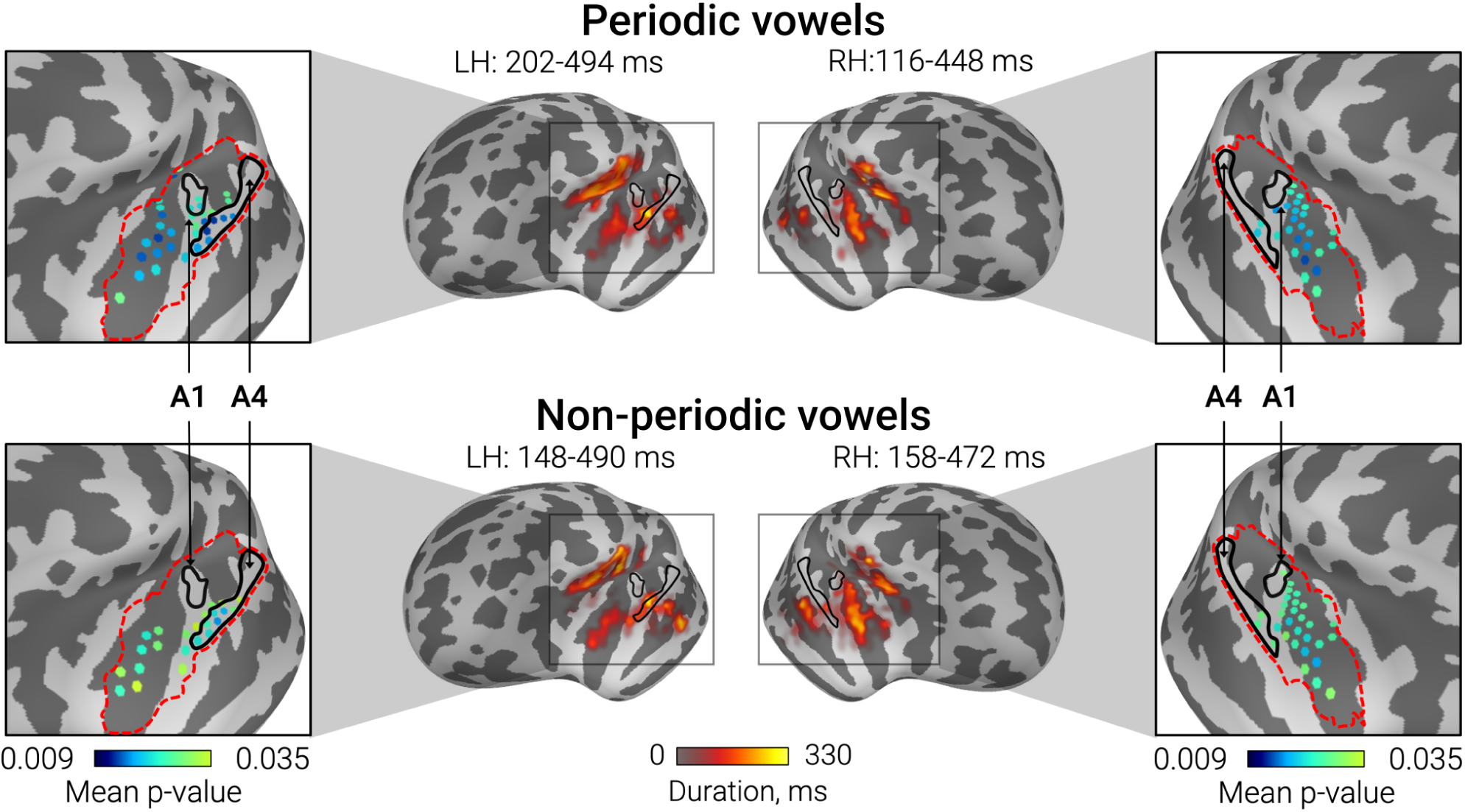
Differences between TD and ASD groups in sustained processing negativity (SPN) associated with periodic and non-periodic vowels. Central panels show the cortical localization of the TFCE clusters of significant group differences in differential responses (vowels minus control non-periodic non-vowel stimulus). The Colored bar below the inflated surfaces indicates temporal extent for sources belonging to the TFCE cluster. The left and right panels show the “most significant” dipole sources selected based on the probability of the SPN group differences within the STG+ region delineated by the red dashed line. Color bars below the images indicate vertices’ p-values averaged over the temporal extent of the cluster.

In all cases, group differences were driven by greater negativity in TD compared to ASD group. As shown in Figure 5, the cortical localization of clusters of significant group differences was remarkably similar for periodic and non-periodic vowels. Within the STG+ region, vertex sources with the highest significance and greatest temporal extent of the differences were located anteriorly and laterally or medially relative to the primary auditory cortex (area A1). In the left hemisphere, the most significant and long-lasting differences were observed in the anterior part of the parabelt auditory area A4 [73]. In the right hemisphere, the differences were most prominent in the posterior segment of the circular insula sulcus (pINS). Additionally, the group differences in SPN were localized to the sources in the superior segment of the circular insula sulcus (sINS) and STS. However, the direction of the current in these sources (Figure S3) as well as their position directly above and below the STG+ sources (see Figure S6) suggest that the group differences in sINS and STS are likely the result of point spread from the auditory cortical areas [74,75].

To sum up, neural processing of sounds with vowel frequency composition differed between children with ASD and TD children in the non-primary auditory cortex of both hemispheres. These group differences in the SPN had a more restricted localization than the SPN itself (see Figure 4). SPN associated with periodicity of non-vocal sound did not differ between the groups.

#### Current timecourses associated with processing of vowel formant structure: SPN and P3a-like response

The analysis of timecourses in the “most significant” sources identified by cluster analysis has shown that group differences emerged starting from ∼ 150 - 200 ms after stimulus onset, and persisted for approximately 200 ms (Figure 6. E, F).

**Figure 6.**
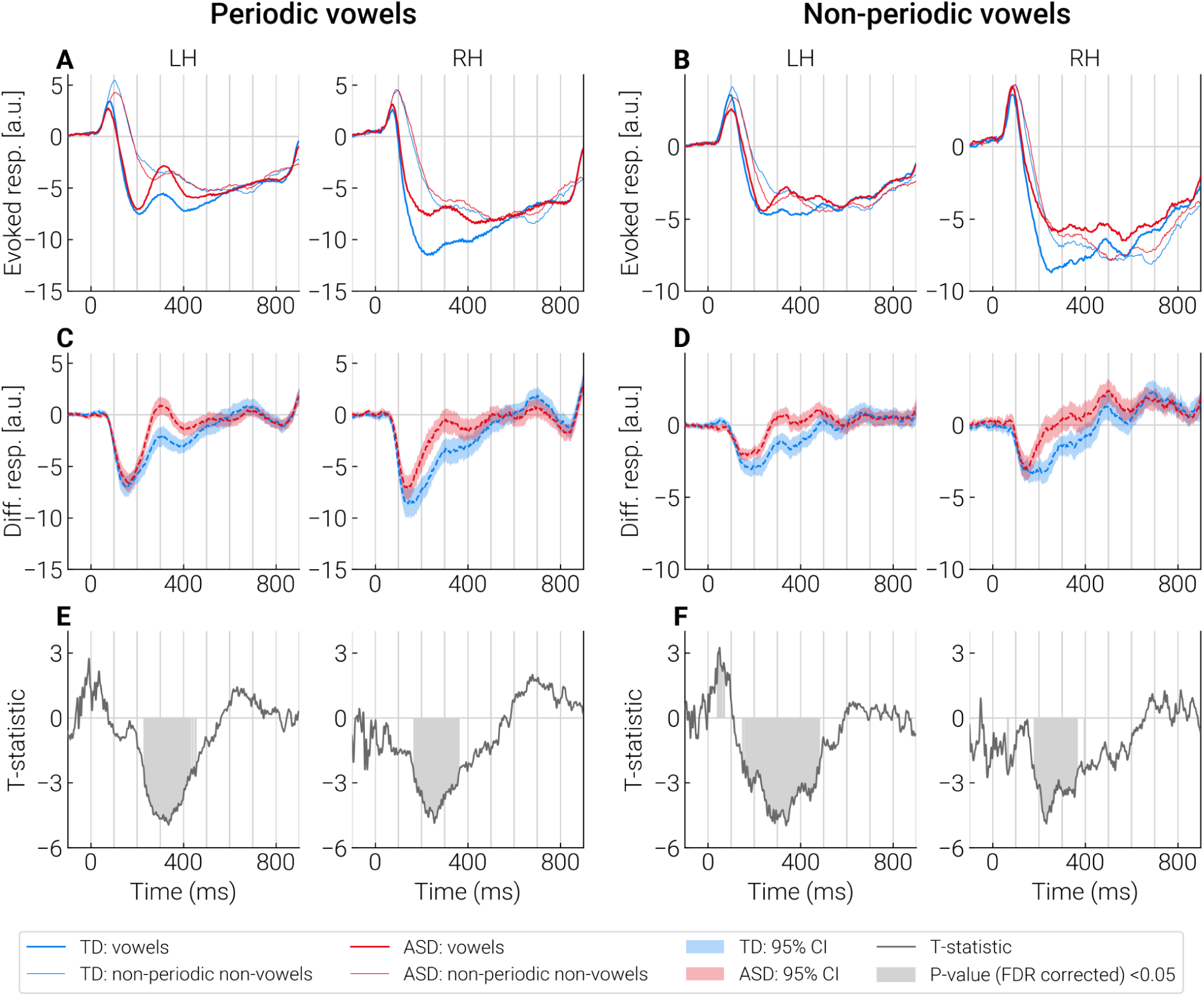
Timecourses of the group differences in differential responses to vowels in the ’‘most significant” dipole sources identified by cluster analysis. **A, B:** Group average timecourses of responses to vowels and control stimuli in TD and ASD groups. **C, D:** Average timecourses of the differential responses (test stimulus - control stimulus) in TD and ASD groups. **E, F:** Point-by-point comparison of timecourses of differential responses between the groups. Significant differences (p < 0.05, FDR corrected) are marked in gray.

To ensure that the differences in the SPN between the groups were due to test rather than control stimuli, we averaged these responses across the “most significant” sources in the interval bounded by the mean onset and end times of significant group differences (see Methods and Supplementary Figure S2). No group differences were found in responses to control stimuli (all p’s > 0.3). The responses to periodic and non-periodic vowels were reduced in children with ASD compared to TD children (periodic vowel, left hemisphere: t(72) = 3.02 p = 0.003, Cohen’s *d* = 0.67, right hemisphere: t(72) = 4.44, p = 0.00003, Cohen’s *d* = 0.92; non-periodic, left hemisphere: t(72) = 1.92, p = 0.06, Cohen’s *d* = 0.44, right hemisphere: t(72) = 3.59, p = 0.0006, Cohen’s *d* = 0.78).

Inspection of Figure 6 revealed a positive transient peak with a latency of ∼300 ms that was most prominent in response to periodic vowels (and less so to non-periodic vowels) in the left-hemisphere. Considering recent intracranial evidence for left-hemispheric predominance of the P3a response associated with processing of novel and potentially salient speech sounds [89], this deflection might reflect an involuntary capture of attention to perceptually salient vowels presented in a sequence of less salient stimuli.

Group differences in this P3a-like response might obscure (or produce) group differences in SPN. To test this possibility, we performed an additional analysis. For each subject, we low-passed the timecourse signals at 10 Hz and estimated P3a amplitude as the amplitude of the largest positive peak in the 200-400 ms range relative to the mean of the two nearest negative peaks. If a positive peak between 200 and 400 ms was absent or indistinct (i.e., its amplitude relative to the preceding or following negative peaks, was below the RMS of the signal amplitude in the baseline period), the P3a-like peak was considered absent and its amplitude was set to zero.

In the pooled sample of participants, the P3a in response to non-periodic vowels was more frequently detected in the left than in the right hemisphere (69% and 50% respectively; Chi square = 4.9, p = 0.02). In the left hemisphere, it was more frequently detected in response to nonperiodic vowels than control stimuli (69% and 45% respectively; Chi square = 8.64, p = 0.003), while no such difference was found in the right hemisphere (50% vs 42%; Chi square = 0.95, p = 0.33). There were no group differences in the occurrence of P3a in response to non-periodic vowels either in the left (TD: 64%, ASD: 74%; Chi square = 0.85, p = 0.36) or right (TD: 44%, ASD: 57%; Chi square = 1.2, p = 0.27) hemispheres, and its amplitude did not differ between the groups (Mann-Whitney U-test, left hemisphere: Z = 1.53, p = 0.12, η2 = 0.03; right hemisphere: Z = 1.29, p = 0.17, η2 = 0.02).

In response to periodic vowels, the P3a-like peak also occurred more frequently in the left than in the right hemisphere in the combined sample of participants (82% and 66% respectively; Chi square = 4.89, p = 0.03) and was more often present in response to periodic vowels than control stimuli (left: 82% and 46% respectively; Chi square = 0.85, p < 0.0001; right: 66% and 43%; Chi square = 7.8, p = 0.005). In response to periodic vowels, P3a in the left hemisphere was more frequently detected in the ASD than in TD group (TD: 72%, ASD: 94%; Chi square = 6.07, p = 0.014), whereas no group differences were found in the right hemisphere (TD: 67%, ASD: 66%; Chi square = 0.01, p = 0.93). Amplitude of the P3a in response to periodic vowels was higher in children with ASD than in TD children in the left hemisphere (Mann-Whitney U-test: Z = 2.93, p = 0.003, η2 = 0.12), but not in the right one (Mann-Whitney U-test: Z = 0.67, p = 0.50, η2 = 0.01).

The amplitude of the left-hemispheric P3a-like response to periodic vowels decreased with age in participants with ASD (Spearman R = - 0.45, p = 0.007), but not in TD participants (Spearman R = - 0.11, p = 0.52), although group differences in correlation coefficients did not reach the level of significance (Fisher’s Z = 1.54, p = 0.13). No correlations with age were found for the left-hemispheric P3a-like responses to non-periodic vowels (TD: R = - 0.12, ASD: R = 0.03, n.s.).

To sum up, the SPN to vowel formant structure was reduced in children with ASD in the STG areas anterior and lateral to the A1 and in pINS. This reduction was observed in both hemispheres from ∼ 150 - 200 ms to ∼ 350 - 450 ms after stimulus onset. In the left hemisphere, periodic and non-periodic vowels evoked P3a-like deflection that was superimposed on the sustained negativity. In response to periodic vowels, the left-hemispheric P3a was present more frequently and with larger amplitude in the ASD group than in the TD group, whereas no group difference was found for the non-periodic vowels. These findings indicated that the P3a-like response might contribute to diminished SPN to periodic vowels in children with ASD in the left hemisphere, but was unlikely to account for the significant reduction of the SPN associated with processing of non-periodic vowels. In the right-hemisphere, P3a-like response can hardly explain the group differences in SPN to either periodic or non-periodic vowels.

### SPN to vowels predicts WiN scores in children with ASD

To test whether atypical SPN to vowels in children with ASD predicts their WiN scores, we applied partial correlation analysis, controlling for age and IQ. SPN was preliminary adjusted (SPNadj) for the magnitude of response to control stimulus and for the number of averaged trials in the following two steps. *First,* for each subject, the timecourses of responses to vowel and control stimuli were averaged over the “most significant” sources in the time interval bounded by the mean start and end times of significant group differences (see Methods and Supplementary Figure S2). *Second*, we computed SPNadj as regression residuals after partialling out the magnitude of response to control stimulus and the square root of the number of averaged epochs from the magnitude of response to the test stimulus.

The partial correlations of SPNadj with WiN scores are presented in Table 2, separately for periodic and non-periodic vowels. For both types of vowels, greater SPNadj in the left hemisphere was associated with better WiN scores in the AM noise (WiNam). Figure 7 illustrates the relationships between adjusted SPN to periodic and non-periodic vowels in the left hemisphere and WiNam performance in the ASD sample. No significant correlations were found for the WiNam in the right hemisphere or for the WiNst in either hemisphere. We also checked for the presence of the partial correlations between SPNadj and WiN scores in TD children. None of the correlations were significant (all p > 0.7).

**Figure 7.**
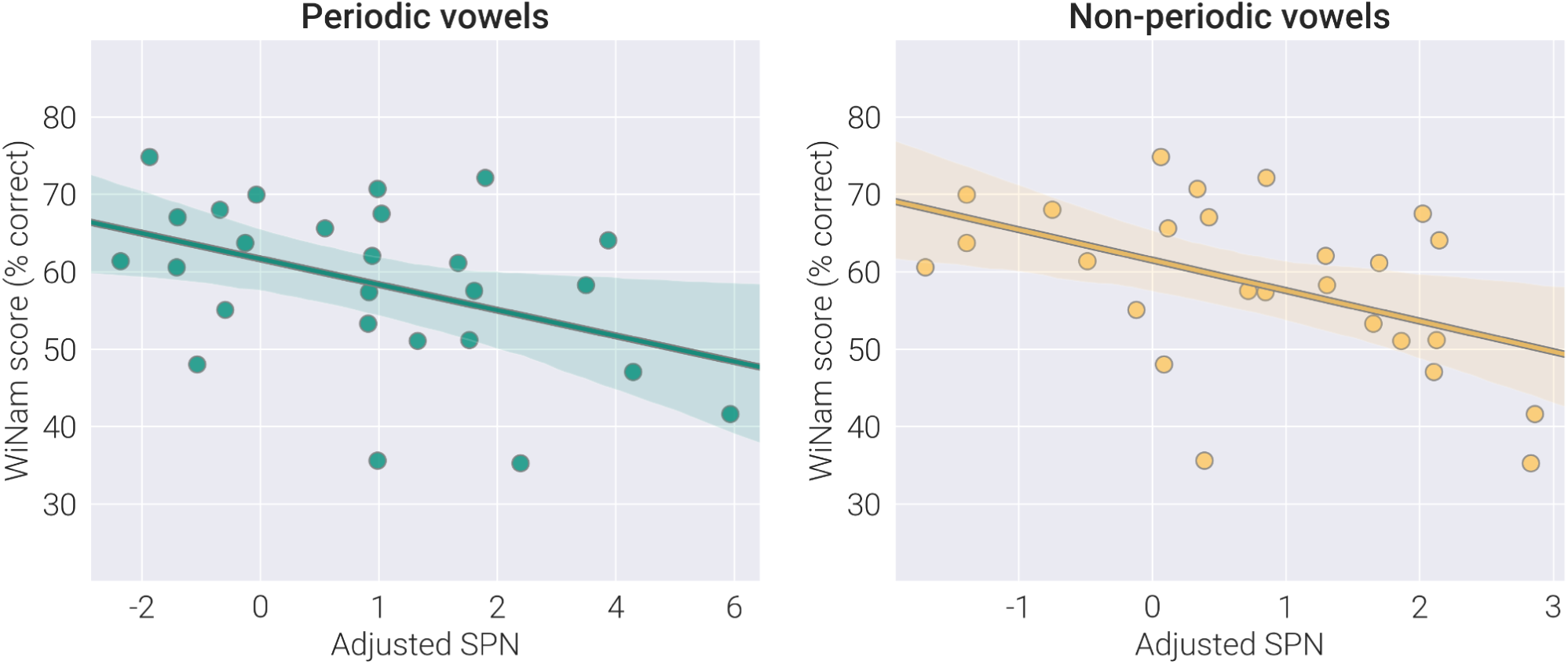
The relationship between WiNam scores and adjusted SPN to vowel stimuli in children with ASD. WiNam scores - percent of correctly repeated words in the amplitude modulated noise. Adjusted SPN - magnitude of the sustained processing negativity adjusted for the magnitude of response to control stimuli and for the number of averaged epochs. WiNam scores were corrected for age.

**Table 2.** Relationship between psychophysical and MEG results in children with ASD: Spearman partial correlation between WiN scores and SPNadj to periodic and non-periodic vowels.

|  | Periodic vowels | Non-periodic vowels |
| --- | --- | --- |
|  | Left hemisphere |  |
| **WiNam** | **SPNadj: $R_{part} = -0.55$, $p = 0.005^*$**<br>Age: $R_{part} = 0.50$, $p = 0.013^*$<br>IQ: $R_{part} = 0.19$, $p = 0.36$ | **SPNadj: $R_{part} = -0.61$, $p = 0.001^*$**<br>Age: $R_{part} = 0.48$, $p = 0.02^*$<br>IQ: $R_{part} = 0.13$, $p = 0.53$ |
| **WiNst** | SPNadj: $R_{part} = -0.26$, $p = 0.22$<br>Age: $R_{part} = 0.46$, $p = 0.02^*$<br>IQ: $R_{part} = -0.21$, $p = 0.32$ | SPNadj: $R_{part} = -0.19$, $p = 0.37$<br>Age: $R_{part} = 0.45$, $p = 0.02^*$<br>IQ: $R_{part} = -0.23$, $p = 0.27$ |
|  | Right hemisphere |  |
| **WiNam** | SPNadj: $R_{part} = -0.03$, $p = 0.88$<br>Age: $R_{part} = 0.52$, $p = 0.01^*$<br>IQ: $R_{part} = 0.10$, $p = 0.64$ | SPNadj: $R_{part} = -0.01$, $p = 0.96$<br>Age: $R_{part} = 0.51$, $p = 0.01^*$<br>IQ: $R_{part} = 0.11$, $p = 0.62$ |
| **WiNst** | SPNadj: $R = -0.02$, $p = 0.92$<br>Age: $R_{part} = 0.49$, $p = 0.01^*$<br>IQ: $R_{part} = -0.23$, $p = 0.28$ | SPNadj: $R = -0.02$, $p = 0.91$<br>Age: $R_{part} = 0.49$, $p = 0.01^*$<br>IQ: $R_{part} = -0.23$, $p = 0.28$ |

WiN - words-in-noise. SPNadj - Sound Processing Negativity, adjusted for response to control stimuli and for the number of averaged epochs. The analysis was performed in 26 children with ASD in whom WiN + IQ + MEG data were available. Significant Spearman partial correlations of interest which survive Bonferroni correction for multiple comparison (uncorrected p < 0.00625) are marked in bold. P-values are two-tailed, uncorrected for multiple comparisons. * p < 0.05 To assess the specificity of the correlations for WiNam, we compared coefficients of correlations of the left-hemispheric SPNadj with the WiNam and WiNst scores using Fisher’s Z statistics for dependent samples [90]. In the case of non-periodic vowels, the correlation was significantly higher for WiNam than WiNst scores (N = 26, Z = 2.35; p = 0.018). No significant difference was found for the periodic vowels (N = 26, Z = 1.60; p = 0.11).

To test whether P3a-like response contributed to the correlation with the WiNam scores we repeated the partial correlation analysis including P3a amplitude as an independent variable (Supplementary material, Table S2). The results suggest that the link between SPNadj and WiNam scores cannot be explained by the differences in the amplitude of the P3a-like component.

Children in the ASD sample had variable levels of IQ, which ranged from moderate intellectual disability to age-appropriate intellectual capacities (Table 1). Therefore, to evaluate the nonspecific effect of disorder severity, we tested whether the reduced SPNadj to vowels in children with ASD was related to their lower IQ scores. No significant Spearman correlations were found (periodic vowel, left hemisphere: R = - 0.03, n.s.; periodic vowel, right hemisphere: R = - 0.21, n.s.; non-periodic vowel, left hemisphere: R = 0.01, n.s.; non-periodic vowel, right hemisphere: R= - 0.2, n.s.).

## DISCUSSION

We investigated whether cortical processing of isolated sounds characterized by vowel formant structure and/or periodicity (pitch) differs in the auditory cortex of children with ASD and TD children, and whether differences related to the processing of these key vowel features contribute to poor perception of words in noise in children with ASD. In both groups of children presence of periodicity and formant structure was associated with sustained processing negativity (SPN) – an early (starting before 100 ms) negative shift of current in the primary and non-primary auditory cortical areas that persisted for several hundred milliseconds. We found no evidence for atypical processing of periodicity (*f0*) of non-vocal spectrally complex sounds lacking formant composition in children with ASD. In contrast, the SPN evoked by vowel-like sounds characterized by formant structure, was significantly reduced in ASD as compared with TD children, regardless of the periodicity of the sound. This SPN reduction emerged relatively late (around 150 - 200 ms after a vowel onset) and was localized bilaterally to the auditory areas anterior to the primary auditory cortex (parabelt area A4 and/or the pINS cortex). In the left, but not in the right hemisphere, reduced SPN in response to vowels predicted poor recognition of words presented against AM noise. Overall, our results suggest that impaired processing of vowel formant composition in children with ASD contributes to their impaired ability to recover words from glimpses of speech interrupted by masking noise.

### Attenuated processing of vowel formant structure in children with ASD

The presence of SPN in response to periodicity or formant structure is consistent with the previous findings in neurotypical children and adults [39,41,88] and extends these findings to children with ASD, at least those with phrasal speech. Similar sustained negative shifts of current were observed in several recent MEG and EEG studies in response to sounds that can be characterized as acoustic patterns distinguished on the basis of their temporal properties such as periodicity [91] or frequency composition, either static or coherently changing in time [30,38,40,92–94]. It has been suggested that this negative shift reflects the fundamental cortical mechanism of automatic grouping in the auditory modality [94]. The early (< 100 ms) latency of the SPN in our study (Figure 4) is consistent with evidence on the remarkably early sensitivity of human auditory cortex to acoustic patterns [30] and, in particular, to vowels [88].

Being well-recognized vocal sounds deeply shaped by the experience of verbal communication, vowels are unique auditory objects. Studies have repeatedly shown that certain areas of the secondary auditory cortex and adjacent regions show a preference for conspecific vocalizations in humans [95–97] and non-human primates [79]. Although still debated [96], it has been suggested that voices are similar to faces in many ways, as both are “special”, carry information about the personality and emotional state of the subject, and are processed in specialized cortical areas [95].

In this respect, the decrease of SPN evoked by periodic and non-periodic vowel-like sounds in children with ASD is a remarkable finding. This decrease can be attributed specifically to an attenuated response to formant composition rather than to the periodicity of the vowel amplitude envelope (fundamental frequency / pitch), as the latter auditory cue was absent in non-periodic vowels. Since children with ASD had normal SPNs to nonvocal sounds characterized by *f0* periodicity, as well as normal responses to control nonperiodic nonvocal stimuli (see Figure 6 A and B), the reduced SPNs to vowels cannot be explained by a general decrease in response amplitude or non-specific deficit in auditory pattern processing.

Despite the early start of the SPN (< 100 ms post-stimulus onset), its group differences emerged relatively late (> 150 ms post-stimulus onset) and were located predominantly in the non-primary auditory areas (Figure 5). Spared functional activity of the primary auditory cortex in children with ASD in our study is in line with fMRI findings in ASD individuals [98,99]. This result is also consistent with the results of Engineer et al. [100] who found in a mice model of autism that non-primary auditory cortical areas are more vulnerable to prenatal factors leading to autism than the primary auditory cortex.

Presence of typical SPN up to at least 150 ms after vowel onset (Figure 6) suggests that the reduced activity in response to vowels in children with ASD is not inherited from the earlier stages of analysis, such as tonotopic processing of formant frequencies [101], detection of harmonicity in complex sounds [102] or detection of an acoustic pattern [30]. On the other hand, timing of the vowel-related SPN reduction generally agrees with results of the meta-analysis of MMN/MMF studies which concluded that responses to phoneme changes (either vowels or syllables) are reduced in individuals with ASD [48]. These considerations suggest that the processing deficit in children with ASD arises at the stage of phonetic analysis.

Notably, the time at which we observed a decrease in SPN in children with ASD coincides with the time at which categorization of isolated vowel sounds into distinct phoneme categories (e.g., /u/ vs. /a/) occurs (∼175 ms post-stimulus onset) [103]. This stage is referred to as acoustic-phonetic mapping and distinguishes brain responses reflecting the true internalized percept of a vowel category from those that index acoustic properties of the vowel [103]. Deficits in phoneme category perception (e.g., relating vowels in the /y/ - /i/ continuum to the category /y/ or /i/) have previously been reported in children with ASD, despite preserved or even superior phoneme discrimination abilities (e.g., judging two vowels in the /y/ - /i/ continuum as the same or different) [104]. Therefore, it is likely that the neurofunctional abnormalities leading to decreased SPN in response to vowels in children with ASD reflect impaired acoustic-phonetic mapping necessary for phoneme categorization. This hypothesis is consistent with the observation that, in the left hemisphere, SPN reduction was most reliable and persistent in the mid-STG region located lateral to the primary auditory cortex in Heschl’s gyrus (parabelt area A4 according to HCPMMP1 [73]) (Figure 5). This area was suggested to be an initial STG waypoint of the ventral auditory stream – the auditory pathway optimized for recognition of acoustic pattern [34,105,106], especially those representing conspecific communication calls [107,108]. In humans, this parabelt auditory region plays a crucial role in phoneme encoding [34,109,110]. The meta-analysis of neuroimaging studies of speech processing [34] concluded that phoneme recognition is associated with activation in the left mid-STG region, while the integration of phonemes into more complex patterns (i.e., words) is localized to the left anterior STG. This conclusion received strong support in a recent study of a patient with extensive lesions of the bilateral STS and left anterior STG, which showed that the intact region of the mid-STG alone can effectively subserve explicit vowel categorization despite the presence of “pure word deafness” [111].

The SPN in children with ASD was also decreased in the temporo-insular regions medially adjacent to the Hershel gyrus and extending in the anterior direction (Figure 5). The pINS has strong structural and functional connections with the auditory cortex [77,112–115] and is responsive to a wide variety of acoustic stimuli [78]. Yet, registration of its neuronal responses in humans [116] and non-human primates [79] has shown that this auditory region of the pINS is sensitive to conspecific vocalizations and can transmit respective auditory information further down to the anterior insula, which is involved in the evaluation of affective signals conveyed by vocal sounds. In the future, it would be interesting to test whether reduced SPN in pINS regions is associated with impaired human voice emotion recognition in ASD [117].

Apart from signals of auditory modality, the pINS comprises neuronal representations of somatosensory, motor, visual, vestibular, limbic signals and is thought to be involved in multisensory integration [118]. The right insula seems to be particularly important for the audiovisual integration [119]. In this regard, it is interesting that the reduction in the SPN induced by vowels in our participants with ASD was strongest in the right pINS (Figure 5). In the future, it is interesting to investigate whether atypical activity or connectivity of the right pINS contribute to severe deficit in audiovisual integration during phoneme recognition in children with ASD [120].

However, it should be noted that the effects found in the insula in our study should be interpreted with caution because the MEG localization error is increased in deep structures such as the insula [74].

### Suppressed processing of vowel formant structure is associated with words in noise perception difficulties in children with ASD

The reduced negativity underlying processing of formant structure in children with ASD predicted the severity of their word recognition problems in the AM noise: the diminished SPN responses to vowels in the left hemisphere were associated with lower WiNam scores (Table 2, Figure 7). This finding has several important implications for interpreting vowel processing deficit and its impact on auditory speech recognition in ASD.

*First,* while WiN performance in children with ASD showed some developmental improvement throughout childhood, neither child’s age nor IQ could explain correlations between the reduced SPN and lowered WiNam scores (Table 2). The lack of correlation between WiN scores and IQ agrees well with the previous findings on the presence of speech-in-noise recognition difficulties even in high-functioning individuals with ASD [13]. On the other hand, our results suggest that these problems may be caused, at least in part, by a deficient vowel processing at the level of the non-primary auditory cortex.

The passive presentation of auditory stimuli and the presence of the SPN deficit at already ∼ 150 - 200 ms after sound onset - i.e., at the preattentive stage of processing - make the potential contribution of higher-order factors such as voluntary attention or motivation unlikely. Yet, involuntary orienting of attention to auditory stimuli may still influence differences between ASD and TD groups. Indeed, the P3a-like responses to periodic and nonperiodic vowels were observed in both TD and ASD children in our study, likely reflecting involuntary shift of attention to perceptually salient speech stimuli [121]. These responses were left-lateralized, consistent with the left-hemispheric bias of the P3a novelty response to speech revealed in the auditory cortex during intracranial recordings in patients [89]. The presence of elevated P3a to periodic vowels in the left hemisphere in our participants with ASD suggests that their reduced negative responses to vowels is unlikely to be due to inattention to the auditory stream containing speech sounds as it was previously suggested [122]. On the contrary, their involuntary attention seems to be captured by perceptually salient periodic vowel stimuli to a greater degree than in TD children.

The excessive P3a-like response could contribute to the decrease in the left-hemispheric SPN to periodic vowels and its correlation with WiNam scores in children with ASD, but it can hardly explain the general trend toward SPN reduction or a common correlation pattern for both periodic and non-periodic vowels. There are several arguments in support of this assumption. *(1)* No group differences in the P3a-like responses or distinct P3a-like peaks were observed in the right hemisphere, despite the prominent right-hemispheric SPN attenuation in ASD vs TD group (Figure 4, 6). *(2)* In case of non-periodic vowels, the group differences in vowel-related negativity started already around 150 ms (Figure 6D), i.e. in the time interval when P3a is not yet evident. *(3)* The group differences in P3a amplitude and in frequency-of-occurrence of the P3a peak were found for periodic vowels only, while in ASD group, the SPN was reduced for both periodic and nonperiodic vowels. *(4)* In the left hemisphere, SPN was a better predictor of WiNam scores in children with ASD than the amplitude of the P3a-like component (Supplementary Table S2).

Although beyond the scope of this paper, the possible role of an enhanced left-hemispheric P3a-like response to speech sounds in autism deserves mention. The previous studies have shown that the P3a can be relatively independent of antecedent negativity. For example, Torppa et al. [123] observed in children with cochlear implants and normal hearing smaller MMN but larger P3a in response to speech sounds. Vlaskamp et al. [124] found that tone duration deviants induced smaller MMN, but larger P3a in children with ASD. It has been hypothesized that the larger P3a reflects increased recruitment of neural resources to compensate for less efficient automatic processing of salient sounds that lie outside the current attentional focus [124].

*Second,* despite the presence of altered vowel-evoked SPN in the auditory cortex of both hemispheres (Figure 5), correlations with WiNam scores were only found in the left hemisphere (Table 2), indicating a specific relationship between the functional integrity of the left secondary auditory cortex and the ability to recognize words in fluctuating noise in children with ASD. Our previous study, which used the same stimuli to compare the SPN responses in neurotypical children and adults [41], showed that the left hemispheric asymmetry in vowel-evoked SPN was present in adults but not in children, in whom SPN responses had the equal amplitude in both hemispheres. The correlation between left-hemispheric but not right-hemispheric SPN to vowels and WiNam scores in children with ASD suggests that some degree of left-hemispheric specialization for vowel processing is already present in childhood and possibly increases with age, driven by the need to integrate the encoding of vowel spectral composition with a predominantly left-lateralized language system (see [125] for the concept of graded hemispheric specialization).

The left STG region that most reliably separated between ASD and TD participants in the present study is remarkably similar to the region that appeared to be sensitive to intelligibility of the sentences that, in turn, depends on the slow temporal modulation of the speech signals at the level of syllables (3-4 Hz) [105]. Our results do not exclude a role of the left A4 region in sentence intelligibility, perhaps in the context of the top-down interactions between phonetic and higher-level (lexical, syntactic, working memory, etc.) processes, but suggest that this region is tuned to spectro-temporal composition of vowels and that weakening of this tuning hinders WiNam task performance.

*Third,* while the atypical left-hemispheric processing of vowels in children with ASD correlated with WiNam scores, it did not correlate with WiNst scores (Table 2). This pattern of correlations suggests that the impaired vowel processing interferes with the ability of listeners with ASD to use dips in noise to capture acoustic cues. Psychoacoustic studies have shown that in subjects with normal hearing, information important for word recognition in fluctuating noise is conveyed through both the temporal fine structure (TFS) of vowels, i.e., carrier frequencies of formants, and their common amplitude envelope (*f0* / pitch) [52]. In our study, WiNam scores in children with ASD correlated with SPN evoked by periodic and non-periodic vowels (Table 2). Since both have a formant structure, but nonperiodic vowels lack *f0*, atypical processing of formant frequencies seems to be a crucial factor contributing to the difficulties in perceiving words in AM noise in children with ASD. However, late (> 150 ms post-stimulus onset) occurrence of SPN reduction and its location in non-primary auditory cortex (left mid-STG region) suggests that the poor “dip listening” is due to insufficient grouping of formants into a “vowel object” rather than decoding of TFS separately for each formant frequency. Consistent with this hypothesis, a recent study of older adults found that impaired central grouping of acoustic patterns is a major contributor to their deficits in processing speech in noise [126].

Our study has several limitations. *First*, we restricted the analysis to temporal cortical regions, where the amplitude of response to sound is maximal, whereas important differences in the processing of linguistic stimuli in autism can also be observed outside the auditory cortex, such as in inferior frontal regions [135]. *Second*, we presented vowel stimuli which are very special overlearned conspecific auditory objects. It would be important to clarify whether the ASD-related deficit in SPN is specific to vowels or whether it is also observed for other auditory objects that have constant or coherently changing frequency composition. *Third,* since we used simple words to test tor speech-in-noise processing difficulties in ASD, one should be cautious about generalizing the findings to more complex linguistic constructions such as sentences. In the case of sentences, speech recognition in noise may be supported by prosody, the slow (syllable-rate) envelope of the speech signal [52], and higher-order semantic cues [136] that are absent or less important in the case of isolated words. *Fourth,* we did not control subjects’ attention to the auditory stimuli, so the possibility remains that differences in attention allocation could affect the results. Comparing the responses in passive and active listening paradigms may help to clarify the role of attention in the observed differences in SPN. *Fifth*, only boys participated in this study. There are multiple sex differences in individuals with ASD (time of diagnosis, genetic burden, neurological and cognitive abnormalities) that may influence the variables investigated in this study [137]. Our sample size did not allow us to analyze the effect of gender. Therefore, we decided to limit our sample to males, who constitute the majority of individuals diagnosed with ASD. More research is needed to see if the findings can be extended to girls with ASD.

### Direction for future research

Word recognition in amplitude-modulated noise depends on multiple integrative processes occurring at different levels of the brain hierarchy and involving numerous feedforward, recurrent, and top-down interactions [127,128]. In a highly heterogeneous ASD population, difficulties with speech perception in noise may arise for a variety of reasons that are attributable to impairments at different stages of the auditory pathway or at higher hierarchical levels. Thus, our results indicating a role for impaired processing of vowel formant structure in WiN perception deficits in children with ASD do not exclude the contribution of other factors. In some children with autism, poor WiN recognition may be due to deficits occurring already at the subcortical level [22–24,129,130], as indexed by the frequency following response to speech sounds [131]. In the future, it would be important to investigate whether impairments in the analysis of the temporal fine structure of sound (TFS) [132] in the brainstem, and the deficit in cortical processes leading to the formation of auditory object contribute independently to poor WiN performance in children with ASD. Our findings do not rule out the “cognitive” hypothesis, which, based on behavioral results, attributes poor masking release in individuals with ASD to a weakness of the domain-unspecific mechanisms that integrate glimpsed fragments into meaningful speech [55,56]. However, since our study was not designed to test this hypothesis, additional neuroimaging research is warranted to address this issue.

Impaired speech-in-noise hearing is one of the central symptoms of auditory processing disorder (APD) - difficulties in recognizing and interpreting sounds that result from central auditory nervous system dysfunction [133] and are often seen in children with ASD and other neurodevelopmental disorders [134]. Detecting at what level of speech signal analysis this dysfunction takes place is important for development of effective and personalized intervention for auditory processing abnormalities not only in ASD, but also in other neurodevelopmental disorders. In this respect, our findings contribute to an emerging profile of children with developmental listening difficulties that may be caused by abnormal processing of speech at different levels of the central nervous system.

## CONCLUSION

Our results suggest that a substantial proportion of children with ASD have altered functional integrity of non-primary auditory cortical areas involved in processing of the vowel formant composition. The localization and relatively late occurrence of this deficit suggest that it arises at the stage of integration of individual formants into phonetic objects – vowels – whereas no deficit was found in children with ASD at earlier stages of processing associated with the encoding of individual formants and/or the detection of a frequency pattern. In the left hemisphere, neural deficit in vowel processing was associated with difficulty recovering words from glimpses of speech in fluctuating noise, a problem characteristic of children with autism. Thus, the impaired grouping of acoustic features into phonetic objects may have an adverse effect on the ability to recognize speech in fluctuating noise in children with autism.

## DECLARATIONS

### Ethics approval and consent to participate

The Ethical Committee of the Moscow University of Psychology and Education approved this investigation, and the principles of the Declaration of Helsinki were followed. Guardians of all children gave written informed consent after the experimental procedures had been fully explained. All participants and/or their caregivers were informed about their right to withdraw from the study at any time during testing.

## Acknowledgments

We sincerely thank children and their families for participating in this study. The study was conducted at the unique research facility “Center for Neurocognitive Research (MEG-Center)” of MSUPE.

## Authors’ contributions

Fadeev K.A.: Formal analysis, Data recording, Data curation, Investigation, Visualization, Software, Writing - original draft; Writing - review & editing;

Romero Reyes, I.V.: Formal analysis, Data curation, Investigation, Visualization, Software, Writing - original draft; Writing - review & editing;

Goiaeva D.E.: Data recording, Investigation, Project administration, Writing - review & editing; Obukhova T.S.: Data recording, Investigation, Project administration, Writing - review & editing; Ovsiannikova T.M.: Data recording, Investigation, Project administration, Writing - review & editing; Prokofyev A.O.: Data recording, Investigation, Writing - review & editing;

Rytikova A.M.: Investigation, Writing - review & editing; Novikov A.Y.: Investigation, Writing - review & editing; Kozunov V.V.: Investigation, Writing - review & editing;

Stroganova T.A.: Conceptualization, Methodology, Funding acquisition, Writing - original draft; Writing

- review & editing.

Orekhova E.V.: Conceptualization, Methodology, Formal analysis, Visualization, Funding acquisition, Writing - original draft, Writing - review & editing;

## Consent for publication

All children gave verbal consent to participate in the study and their caregivers gave written consent for publication of anonymized data.

## Data/code availability statement

The de-identified individual-level raw data that supports this research, study materials, and analysis code used to generate the results are publicly available at https://openneuro.org/datasets/ds005234

## List of abbreviations

ASD: autism spectrum disorder
AM: amplitude-modulated [noise]
EEG: electroencephalography
ERP/ERF: event-related potential/field
INS: insula
MEG: magnetoencephalography
MMN/MMF: mismatch negativity/field
MPI: Mental Processing Index
ROI: region of interest pINS - posterior insula
SCQ: Social Communication Questionnaire
sLORETA: standardized low-resolution brain electromagnetic tomography
SNR: signal-to-noise ratio
SPN: sustained processing negativity
SPNadj: SPN adjusted for the response to the test stimulus and for the number of averaged epochs.
SRS: Social Responsiveness Scale
ST: stationary [noise]
STG: superior temporal gyrus
STS: superior temporal sulcus
TD: typically developing
TFCE: Threshold-Free Cluster Enhancement
WiN: words in noise
WiNam: words in amplitude-modulated noise
WiNst: words in stationary noise

