## Supplementary Materials (Method, Figures, Tables) for "Attenuated processing of vowels in the left hemisphere predicts speech-in-noise perception deficit in children with autism"

**to**

**Supplementary Methods**

***I. Assessing the general level of language development***

All test stimuli were presented through the AutoRAT application (Ivanova et al., 2016) installed on a Samsung Galaxy Tab A (2016) SM-T585 running Android 7.0, with a 10.1” screen with a resolution of 1920x1200 pixels. The description of the components of the RuCLAB test (Lopukhina et al., 2019) is given in Table S1. Subjects’ responses were audio-recorded and analyzed offline.

In all language reception tasks, correct answers were coded as "1" and incorrect answers were coded as "0". Incorrect pronunciation of sonorant phonemes, in particular [r], [r'], [l], and [l'], was not considered an error.

In word production tasks, accuracy was also assessed by coding responses as "1" for correct and "0" for incorrect. An answer was considered correct if participants accurately named the depicted object or action (e.g., "ant" for depicted ant) or provided a subdominant nomination. In the sentence composition test, accuracy was evaluated in three parameters: a) morphosyntactic correctness, b) lexical and semantic accuracy, and c) integration of other sentence components. Each correctly completed parameter was scored 1 point. These points were summed up and divided by 3. For example, a sentence that met two of the three criteria would receive 0.66 points (2/3). In the discourse production subtest, accuracy was assessed according to four parameters: a) fluency, b) morphosyntactic correctness, c) content richness, and d) absence of semantic, phonological and other errors. In this subtest, each parameter was scored with a maximum of 5 points. These scores were summed and divided by 20. For example, a total sum of 15 points (5+3+4+3) would give a final test score of 0.75 (15/20).

*References*

Lopukhina, A., Chrabaszcz, A., Khudyakova, M., Korkina, I., Yurchenko, A., & Dragoy, O. (2019). Test for assessment of language development in Russian «KORABLIK». In Proceedings of the Satellite of AMLaP conference «Typical and Atypical Language Development Symposium» (p. 30).

Ivanova, M., Dragoy, O., Akinina, J., Soloukhina, O., Iskra, E., Khudyakova, M., & Akhutina, T. (2016). AutoRAT as your fingertips: Introducing the new Russian Aphasia Test on tablet. Frontiers in Psychology Conference Abstract: 54th Annual Academy of Aphasia Meeting. <https://doi.org/10.3389/conf.fpsyg.2016.68.00116>

### II. Words-in-Noise (WiN) test scoring: criteria for exclusion of the “easiest” and “ most difficult” words from analysis

Although all of the two-syllable words used in the WiN test were frequent and familiar to children with autism, some were particularly easy or difficult to recognize. These words were identified and excluded from the analysis based on the behavioral results from our previous study (Fadeev et al., 2023). This study included a sample of 50 TD and 47 ASD children aged 7-12 years and used the same WiN psychophysical test as the present study. For each word, we estimated the percentage of successful recognitions in each of the 6 conditions (AM: 0, -3, -6 dB SNR and ST: 0, -3, -6 dB SNR), separately in the TD and ASD groups. Then, separately for each condition and group, these percentages were converted to z-scores. The z-scores were averaged across groups and conditions, so that each word was "ranked" in terms of its “ease of recognition”. Words with z-scores above or below one standard deviation from the mean were excluded from the analysis. Thus, we excluded the 8 "easiest" and 10 "most difficult" words.

*References*

Fadeev, K. A., Goyaeva, D. E., Obukhova, T. S., Ovsyannikova, T. M., Shvedovskiy, E. F., Yu. Nikolaeva, A., Davydova, E. Y., Stroganova, T. A. & Orekhova, E. V. (2023). Difficulty with Speech Perception in the Background of Noise in Children with Autism Spectrum Disorders Is Not Related to Their Level of Intelligence. Clinical Psychology and Special Education, 12(1), 180–212. <https://doi.org/10.17759/cpse.2023120108>

**Supplementary Table S1.** Components of the RuCLAB test

| **Test Component** | *Linguistic level* | *Description* |
| --- | --- | --- |
| **Phonological discrimination**  **(receptive test)** | phonology | Two pseudowords were auditorily presented to the children. If the words were the same, they were instructed to press the "YES" button on the tablet screen; if they were different, they were instructed to press the "NO" button.  N = 26* |
| **Nonword repetition**  **(expressive test)** | phonology | Evaluating the capacity to repeat heard non-existing “words”.  N = 26 |
| **Word repetition**  **(expressive test)** | vocabulary | Assessing the ability to repeat heard words.  N = 26 |
| **Object naming**  **(expressive test)** | vocabulary | Testing vocabulary and word retrieval speed.  Participants name visually presented objects.  N = 26 |
| **Action naming**  **(expressive test)** | vocabulary | Assessing verbal fluency and comprehension of actions. Participants describe the actions depicted in the images.  N = 26 |
| **Noun**  **comprehension**  **(receptive test)** | vocabulary | Four black and white images of objects are displayed on the tablet. Upon hearing a noun corresponding to one of the images, the participant must select the correct image by tapping on it on the tablet. The three incorrect images represent semantic, phonetic, and irrelevant errors.  N = 26 |
| **Verb comprehension**  **(receptive test)** | vocabulary | Similar to the noun comprehension subtest, but with verbs.  N = 26 |
| **Sentence repetition**  **(expressive test)** | morphosyntax | Requires listening to sentences and repeating them as accurately as possible. Tests auditory memory, language comprehension and verbal reproduction skills.  N = 14 |
| **Sentence comprehension**  **(receptive test)** | morphosyntax | Requires listening to sentences and then selecting the appropriate picture from the pair. The subtest assesses language comprehension, auditory processing, and the ability to link verbal descriptions to visual representations.  N = 26 |
| **Sentence production**  **(expressive test)** | morphosyntax | Participants must describe a picture in one sentence, incorporating all essential details to convey a comprehensive picture of the scene.  N = 26 |
| **Discourse comprehension**  **(receptive test)** | discourse | Assessment of narrative comprehension, memory for details, and inferential thinking. Participants listen to a story and then answer questions about its events and details.  N = 20 |
| **Discourse production**  **(expressive test)** | discourse | Evaluating creative thinking, narrative skills, and the ability to organize thoughts cohesively. Participants are asked to creating a story from a given picture, structuring it with an introduction, a climax, and a conclusion.  N = 1 |

* N - number of trials in the subtest.

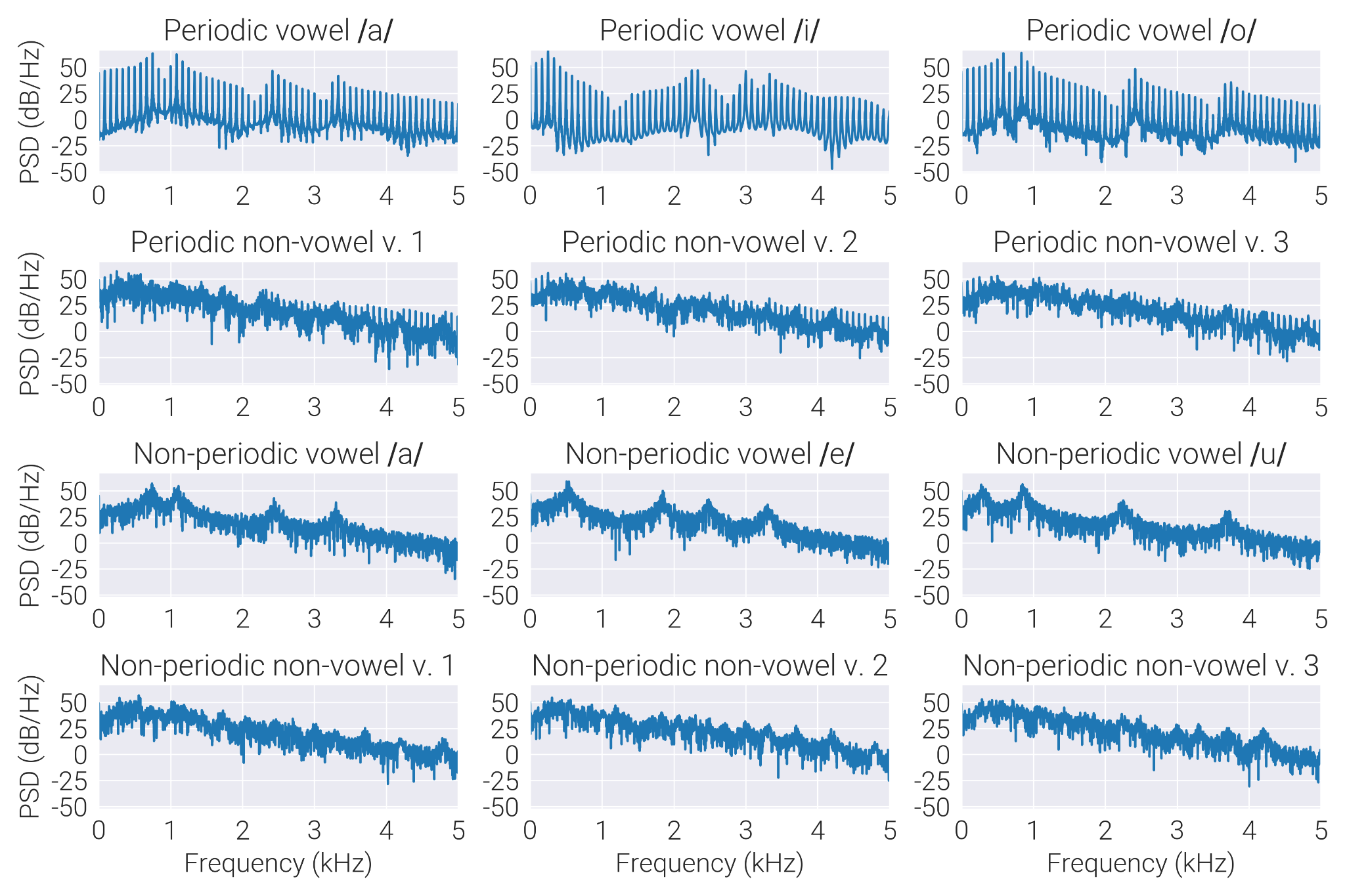

**Supplementary Figure S1** - Spectral composition of four classes of stimuli; each has three variants.

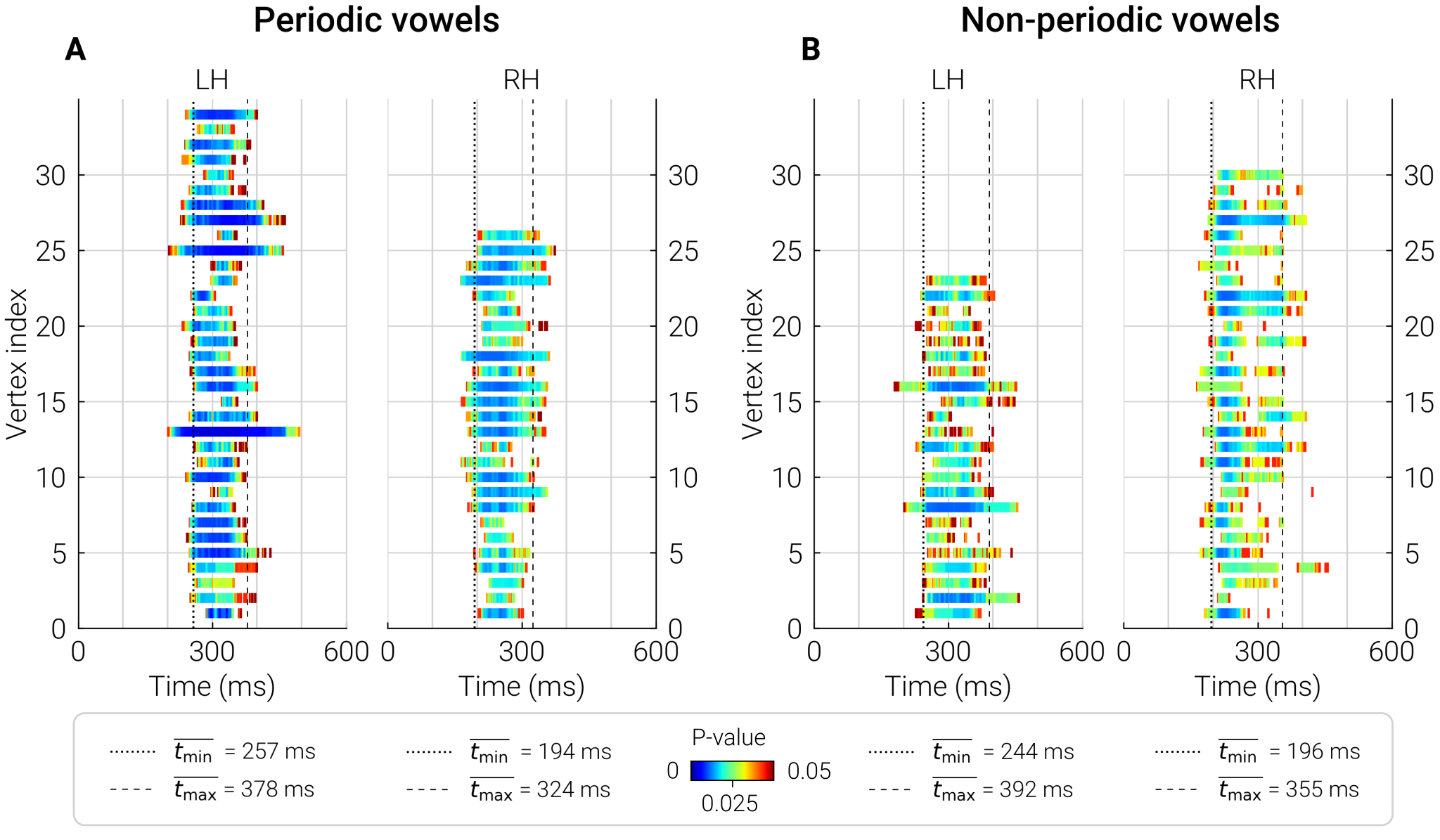

**Supplementary Figure S2** - Time frames of the “most significant” vertices separating between differential responses (i.e., test - control) in the TD and ASD groups. Horizontal bars show time range and probability of the significant group differences in SPN evoked by periodic (A) and non-periodic (B) vowels in each vertex in the “most significant” sources. Average start and end times of these bars are marked with vertical dashed lines and delineate the time region in which the response was averaged.

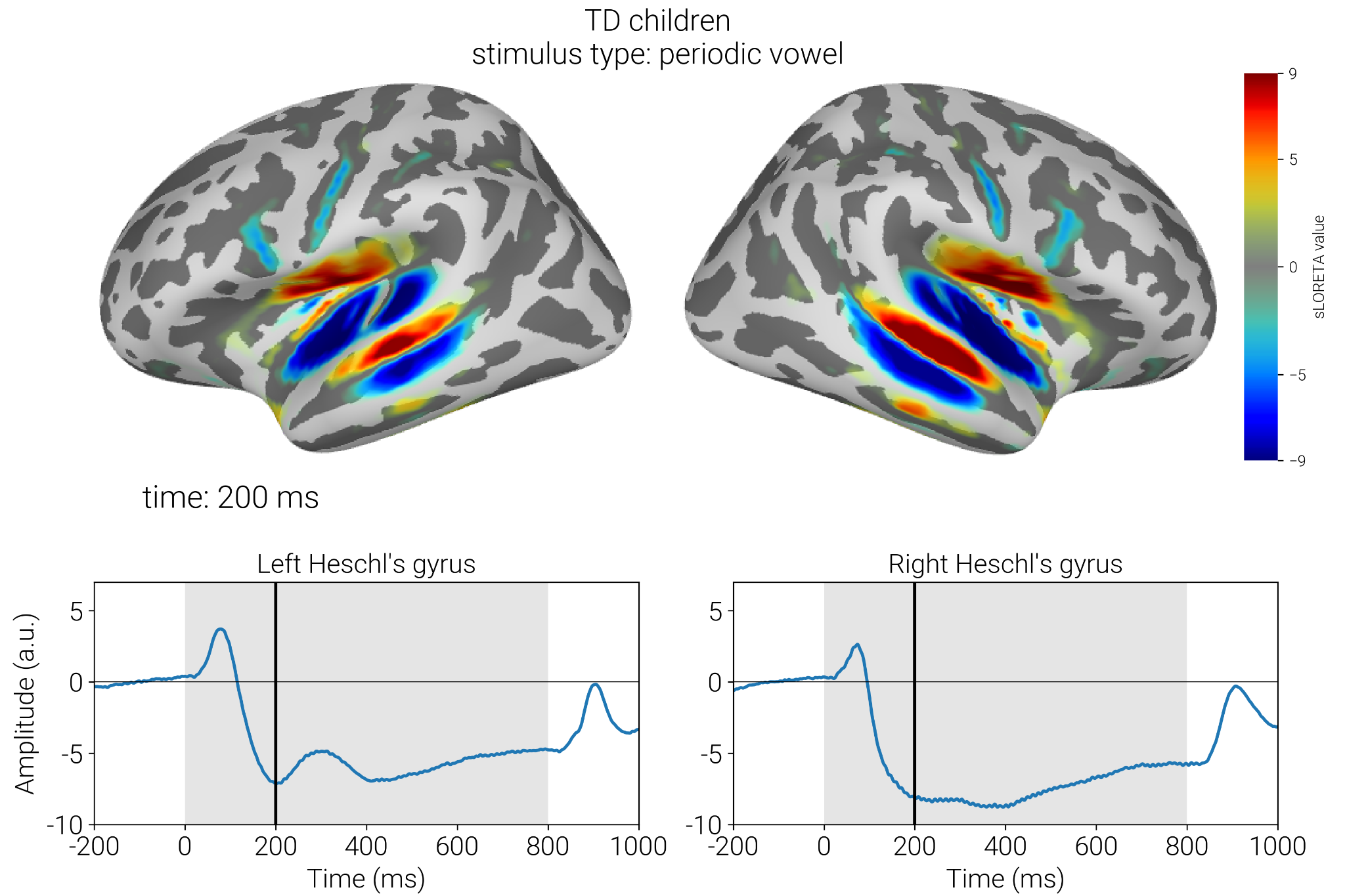

**Supplementary Figure S3** - Grand average response to periodic vowels in TD children. Upper panel shows distribution of sLORETA values at the peak of N2 component; color marks direction of the current. Lower panel show timecourses of the response in the Heschl’s gyri. Changes in the direction of current from STG to superior insula and from STG to STS, suggest that activity observed in these regions is the spread from the STG.

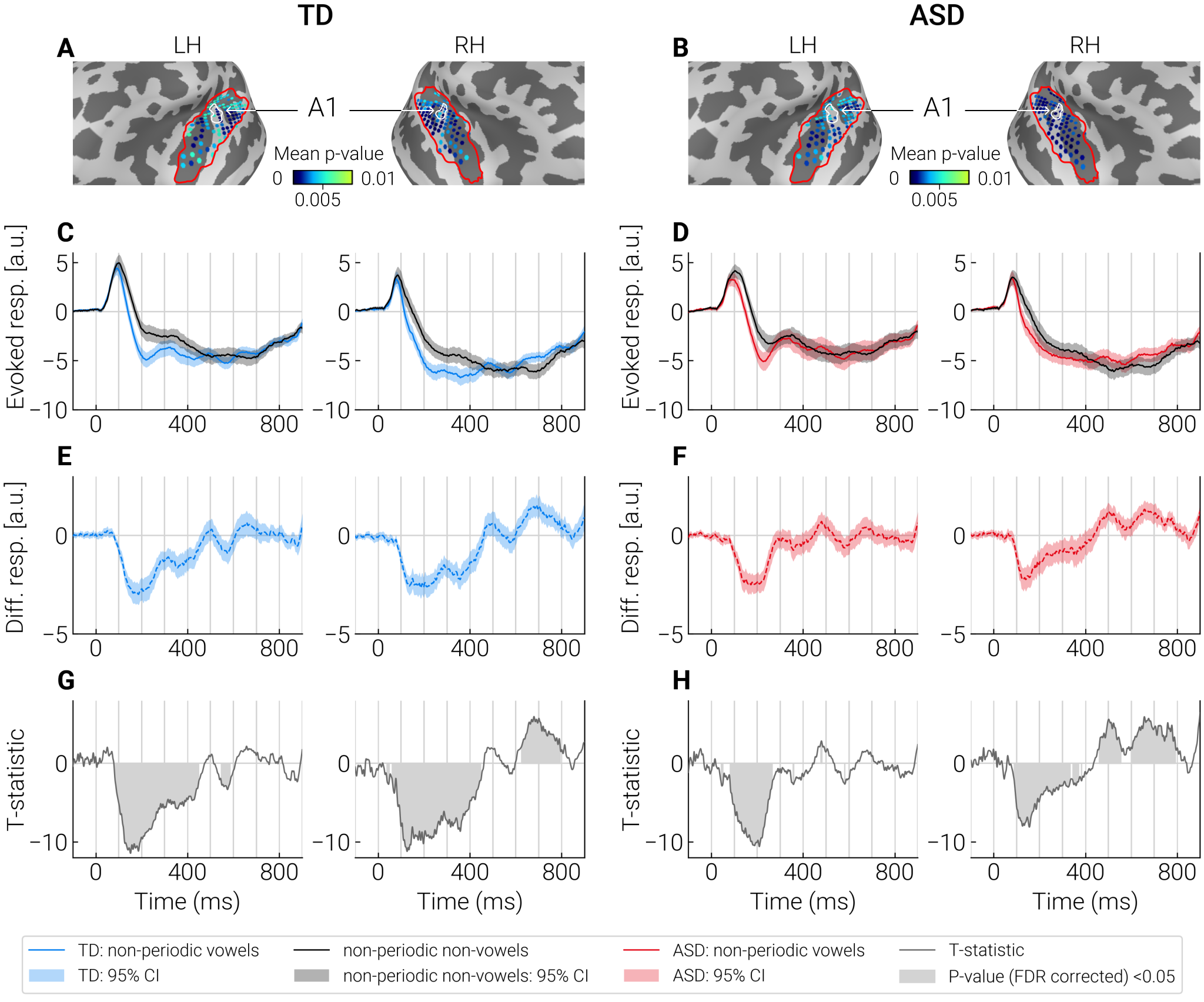

**Supplementary Figure S4** - Comparison of evoked responses to the non-periodic vowels and control stimuli (non-periodic non-vowels) in the “most significant” dipole sources identified by cluster analysis. **A, B:** The “most significant” dipole sources within STG+ region (outlined with a red contour) are marked by blue dots. Color shade (light blue to dark blue) indicates significance of the differences between test and control conditions in the respective point sources. The primary auditory cortex (A1) is outlined with a white contour. **C, D:** Averaged neural current timecourses in the “most significant” sources. **E, F:** The difference between timecourses of current evoked by the test and control stimuli. **G, H:** T-statistics reflecting a pointwise comparison of the response timecourses to test and control stimuli. Significant differences (p < 0.05, FDR corrected) are marked in gray.

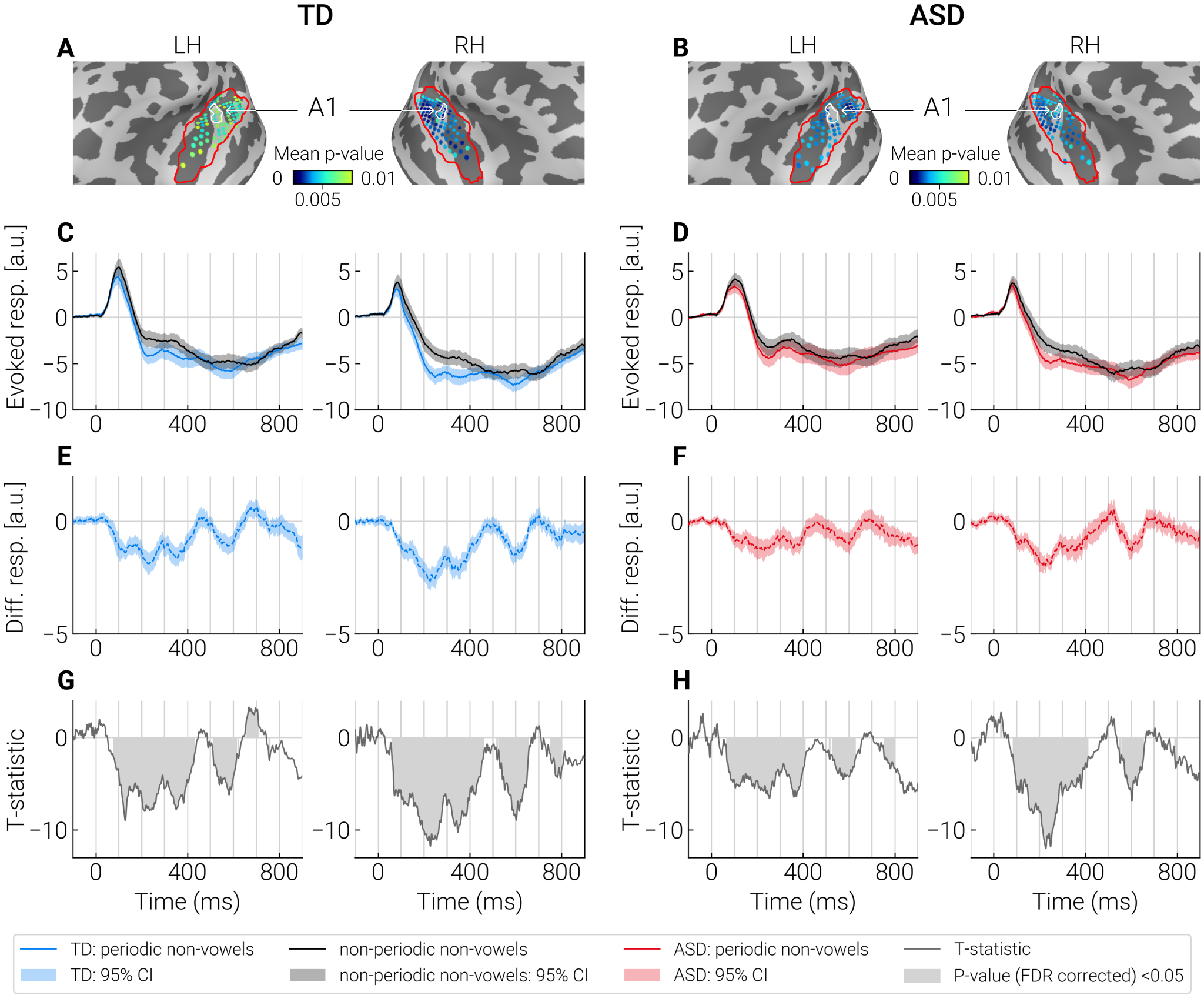

**Supplementary Figure S5** - Comparison of evoked responses to the periodic non-vowels and control stimuli (non-periodic non-vowels) in the “most significant” dipole sources identified by cluster analysis. **A, B:** The “most significant” dipole sources within STG+ region (outlined with a red contour) are marked by blue dots. Color shade (light blue to dark blue) indicates significance of the differences between test and control conditions in the respective point sources. The primary auditory cortex (A1) is outlined with a white contour. **C, D:** Averaged neural current timecourses in the “most significant” sources. **E, F:** The difference between timecourses of current evoked by the test and control stimuli. **G, H:** T-statistics reflecting a pointwise comparison of the response timecourses to test and control stimuli. Significant differences (p < 0.05, FDR corrected) are marked in gray.

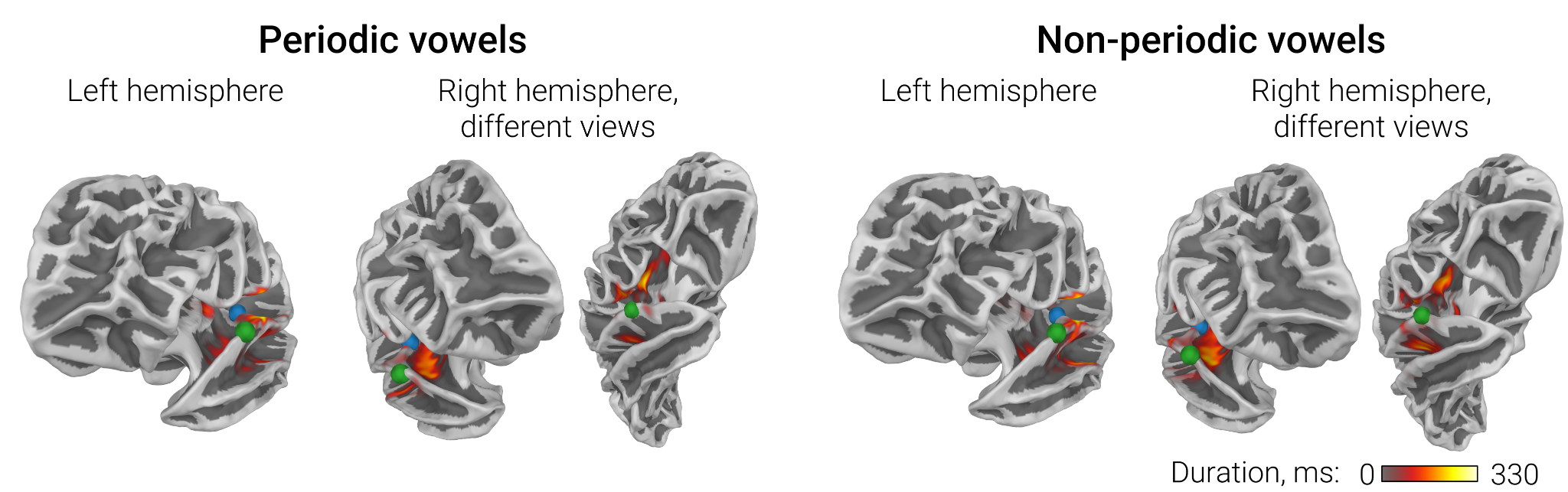

**Supplementary Figure S6** - ASD versus TD group differences in sustained processing negativity (SPN) associated with processing of periodic and non-periodic vowels: results of the TFCE cluster analysis. Clusters of group differences are projected to the white matter surface of the brain. Color scale indicates the temporal extent of the source in the TFC clusters. Blue dots are placed on Heschl's gyri, green dots mark the anterior aspect of the parabelt area A4.

**Supplementary Table S2**. Spearman partial correlation between WiN scores and SPNadj to periodic and non-periodic vowels in the ASD group.

|  | Periodic vowels, left hemisphere | Non-periodic vowels, left hemisphere |
| --- | --- | --- |
| WiNam  (N=26) | **SPNadj: R*part* = -0.50, p =0.015***  Age: R*part* = 0.46, p = 0.03*  IQ: R*part* = 0.17, p = 0.43  P3a: R*part* = -0.06, p = 0.80 | **SPNadj: R*part* = -0.57, p = 0.005***  Age: R*part* = 0.41, p = 0.05  IQ: R*part*= 0.04, p = 0.86  P3a: R*part* = -0.3, p = 0.16 |

**Note.**

P3a - amplitude of the P3a-like component. Correction for P3a amplitude did not significantly affect the correlation between SPNadj and WiNam (see Table 2 in the main text).

**Supplementary Videos S1-6.** Temporal evolution of significant clusters of the differences in evoked source current between test (periodic vowels, non-periodic vowels, periodic non-vowels) and control (non-periodic non-vowels) conditions in TD and ASD groups. Blue color corresponds to SPN, i.e., more negative source current to the test compared to the control condition; red color corresponds to the opposite direction of the difference.

|  | Group | |
| --- | --- | --- |
| Test condition | TD | ASD |
| periodic vowel | **Video S1** | **Video S4** |
| non-periodic vowel | **Video S2** | **Video S5** |
| periodic non-vowel | **Video S3** | **Video S6** |
